# Clonal dynamics of normal hepatocyte expansions in homeostatic human livers and their association with the biliary epithelium

**DOI:** 10.1101/2021.07.02.450704

**Authors:** AM Passman, MJ Haughey, E Carlotti, MJ Williams, B Cereser, ML Lin, S Devkumar, JP Gabrield, FP Russo, M Hoare, J Chin-Aleong, M Jansen, NA Wright, HM Kocher, W Huang, MR Alison, SAC McDonald

**Affiliations:** Centre for Cancer Genomics and Computational Biology, Barts Cancer Institute a CR-UK Centre of Excellence, Queen Mary University of London, London, UK; School of Mathematical Sciences, Queen Mary University of London, London, UK; Centre for Tumour Biology, Barts Cancer Institute a CR-UK Centre of Excellence, Queen Mary University of London, London, UK; Department of Surgery, Oncology and Gastroenterology, University of Padova, Padova, Italy; Cancer Research UK Cambridge Institute, University of Cambridge, Cambridge, UK; Histopathology, Barts Health NHS Trust, London, UK; Department of Cellular Pathology, University College London, London, UK; UCL Cancer Centre, University College London, London, UK; Cancer Tissue Bank, Barts Cancer Institute a CR-UK Centre of Excellence, Queen Mary University of London, London, UK; Barts and the London HPB Centre, The Royal London Hospital, Barts Health NHS Trust, London, UK; Group of Theoretical Biology, The State Key Laboratory of Biocontrol, School of Life Science, Sun Yat-sen University, Guangzhou, China

**Keywords:** Clonal dynamics, stem cell, hepatocytes, biliary epithelium

## Abstract

The majority of human liver research is disease-focused such that far less is known of cellular dynamics within normal human liver. We have leveraged cytochrome *c* oxidase deficiency as a marker of clonal hepatocyte populations in such tissues. We demonstrate these populations commonly associate with portal tracts and lineage-trace hepatocytes with cholangiocytes, indicating the presence of a bipotential common ancestor at this niche. We also observe rare periportal SOX9^+^ hepatocytes progenitor candidates in our human tissues. To understand clonal expansion dynamics, we measured methylation diversity and identified mtDNA variants by next-generation sequencing within spatially-defined clonal hepatocyte patches. We coupled our sequencing with mathematical modelling and Bayesian inference to compare spatial patterns of mtDNA variants under assumptions with or without faster expansion from a portal-associated niche. These datasets support the existence of a periportal progenitor niche and indicate that clonal patches slowly expand, perhaps due to acute environmental stimuli, then quiesce. These findings crucially contribute to our understanding of hepatocyte dynamics in normal human liver and provide a baseline for understanding how such dynamics may be modulated in diseased liver.

## Background

The liver possesses an extensive playbook for cell renewal and recovery from injury. Decades of partial hepatectomy (PH) studies have established that mature hepatocytes, regardless of their anatomic location, are capable of restoring the vast majority of the liver mass (1, 2). Further still, serial PH and hepatocyte transplantation studies have demonstrated that hepatocytes are seemingly inexhaustible in their replicative capacity; all without signs of malignant transformation (3, 4). Hepatocytes are also capable of dedifferentiation, expansion and redifferentiation, both *in vitro* (5) and in rodent damage models (6, 7), and thus may themselves, be capable of regenerating the biliary epithelium (8, 9). Self-renewal and multipotentiality are properties of stem/progenitors and are unusual for a differentiated cell. As such, the necessity for a dedicated stem/progenitor cell and niche, as is present in many other organs, has been called into question.

Regardless of hepatocyte plasticity, isolates of cells variously known as liver progenitor cells, oval cells or hepatic progenitor cells have in fact demonstrated bipotentiality and are thought to reside at a periportal niche (10–12). *In vivo*, contribution of these cells to liver regeneration has been widely reported in rodent injury models (13–15). This has however, been the source of some controversy, with several lineage-tracing studies failing to detect significant proportions of biliary-derived hepatocytes upon regeneration from injury (16, 17). Evidence is now mounting that widespread impairment of hepatocyte replication is required for activation of the progenitor cell regenerative compartment (13, 14, 18). This finding is particularly relevant for human disease in which widespread hepatocyte impairment is commonplace.

Whilst there is a reasonable degree of clarity regarding the cellular responses upon liver injury, there is still conflict regarding homeostatic dynamics. Early studies utilised tritiated thymidine to track the migration of hepatocytes in normal rat livers over time and found a periportal peak of labelled (DNA-synthesising) cells soon after injection (19, 20). Examination of rats sacrificed over time revealed a migration of labelled hepatocytes along the portal-central axis; a phenomenon coined ‘the streaming liver’. The streaming liver hypothesis likened the portal-central hepatocyte chords to intestinal crypts within which, in time, enterocytes progress from the crypt base to villus tip. In the same manner, hepatocytes were proposed to stream from the portal tract (PT) and end their lives at the central vein (CV).

This theory proved contentious (21) and subsequent studies have also proposed neo-hepatocyte generation from all three zones in the hepatic lobule; periportal (22), mid-zonal (17, 23) and pericentral (24). Further still, there is evidence that homeostatic turnover results from zone-agnostic, broadly distributed hepatocytes (25–28).

There are exceptionally few studies of homeostatic dynamics in normal human liver, however there is evidence supporting the existence of a periportal progenitor niche. Huch et al. were able to generate bipotential organoids from EPCAM^+^ ductal cells isolated from normal human livers (29). Similarly, in a single-cell RNAseq atlas of 10,000 normal human liver cells, Aizarani et al. identified a periportal TROP2^int^ progenitor population that was able to form bipotent liver organoids (30). Our approach to studying homeostatic dynamics in human liver centres on demarcation of clonal boundaries within these tissues. In contrast to diseased liver, clones within normal liver are often modest in size (31), likely reflecting low cell turnover. Regardless, clonal patches do indeed exist within normal liver as demonstrated in human lineage tracing studies using cytochrome *c* oxidase (CCO) deficiency as a marker (32, 33). Interestingly, these studies highlighted an association between the clonal patches and the PT. This is intriguing as it is thought that CCO-deficient cells expand from stem/progenitors (33, 34). This is because mutant mitochondria clonally expand by random intracellular drift taking place over decades, and stem/progenitors are long-lived enough to acquire the homoplasmy, or a high enough burden of heteroplasmy (~<75 %) required for enzymatic deficiency (35, 36). Further evidence that the periportal region is a stem/progenitor niche in normal human liver is lacking. Whether potential progenitor cells exist as SOX9^+^ hybrid hepatocytes (37) or as a subpopulation of biliary epithelial cells, is also currently unknown.

In this study we demonstrate for the first time, the spatial dynamics and the origins of human hepatocytes *in vivo*. We utilise a range of techniques including immunohistochemistry, methylation sequencing, mitochondrial DNA next generation sequencing and mathematical modelling to establish their origins and the nature of their expansion. Collectively, our data support the existence of a periportal stem cell niche that feeds the clonal expansion of hepatocyte patches in normal human liver. These patches have a common cell of origin with biliary epithelium and expand in a slow and punctuated fashion, with a portal to central trajectory, and are largely quiescent, potentially for decades thereafter.

## METHODS

### Samples

Human liver samples were obtained by written informed consent from patients undergoing hepatic resection, most commonly for metastatic colorectal cancer. Samples were obtained through the Queen Mary University of London Cancer Tissue Bank (REC reference 14/LO/2031 renewed 19/LO/1700, more information on https://www.cancertissuebank.org, and project approval2015/01/QM/SM/E/FrozenTissue) and University of Cambridge (16/NI/0196). Additional clinical details such as patient age, treatments received and reason for resection, where available, can be found in **Supplementary Table 1**.

### Preparation of tissue

Normal liver was taken from the resection, distal to the metastatic or diseased tissue and normal morphology was identified by a pathologist. Resections were cut to size and normal regions were identified by a pathologist, submerged in isopentane and frozen in liquid nitrogen. Samples were stored at −80 °C until required for cryosectioning. Frozen samples were surrounded with optical cutting temperature medium and sectioned onto UV-treated P.A.L.M. membrane slides (Zeiss, Cat # 415190-9041-000) for laser capture microdissection or charged glass slides for immunohistochemistry.

### Immunohistochemistry

Frozen sections were thawed by air drying, then fixed in 4% paraformaldehyde. Slides were then permeabilized with 0.1% TritonX-100 in tris-buffered saline (TBS) at RT. The tissue was then blocked for endogenous phosphatases and peroxidases with BLOXALL® (Vector Laboratories, Cat # SP-6000-100) or 0.3% H_2_O_2_ before application of 5% horse serum epitope block. The serum was carefully removed, then primary antibody (anti-SOX9 at 1/200 with 0.1% TritonX-100, ABCAM, Cat # ab185966; anti-KRT19 at 1/100, DAKO Cat # M0888) diluted in diluent (DAKO, Cat # S302283) was applied for 1 hr at RT or overnight at 4 °C. Depending on the antibody source and chromogen required, HRP or AP-conjugated Horse Anti-Mouse or Rabbit polymers (Vector Laboratories, Cat # MP-5402 and MP-7401) were applied for 30 min. Chromogen development was then conducted with HRP 3,3’-Diaminobenzidine (DAB) or Vector Blue kits (Vector Laboratories, Cat # SK-4100 or SK-5300). Chromogen development was halted by submersion in tap water, then slides were counterstained with Nuclear Fast Red (Vector Laboratories, Cat # H-3403), cleared with Histoclear (national diagnostics, Cat # HS-200) and mounted with VectaMount^®^, (Vector Laboratories, Cat # H-5000). Double staining was performed sequentially, following the chromogen development and the procedure was repeated beginning at the epitope block.

Anti-Ki67 (1/1000, ABCAM, Cat # ab16667) staining was performed by autostaining on a Discovery Ultra Ventana using the Discovery OmniMap anti-Rb HRP and Discovery ChromoMap DAB kits (Roche, Cat # 760-4311 and 760-159).

### Enzyme histochemistry

A dual-colour enzyme histochemical staining protocol was employed to identify clonal cytochrome *c* oxidase (CCO) deficient patches/cells as previously described (38). This system results in DAB deposition in regions of functional CCO activity. DAB development crowds out secondary deposition of Nitrotetrazolium Blue chloride (NBT), thus by virtue of succinate dehydrogenase (SDH) activity, blue staining occurs only in regions of CCO deficiency. Enzymatic chromogen development of DAB and NBT for CCO and SDH activity were typically halted after 50 and 25 min respectively. We performed CCO/SDH staining in two sets of serial sections. For the first, we cut 6 μm serial cryosections, and utilised serial sections for Ki67 and SOX9^+^KRT19^+^ staining. In the second set, we cut 10 μm serial cryosections, with subsequent serial sections used for laser capture microdissection followed by next generation sequencing. Following sectioning, all slides were stored at −80 °C until use.

### CCO patch-structure association and digital cell counting

Serial sections stained with H&E or for CCO/SDH, SOX9 + KRT19 and Ki67, were scanned with a NanoZoomer S210 (Hamamatsu), then analysed within QuPath (39). All CCO-deficient patches were manually annotated on the CCO/SDH slide. All portal tracts (PT) and central veins (CV) were annotated on H&Es, checked by referencing KRT19 staining, then transferred to the CCO slide. The distances to the nearest CV and PT were then obtained for each CCO patch using the script in Script modified from Pete Bankhead’s post https://forum.image.sc/t/qupath-distance-between-annotations/47960/2. PT, CV and CCO annotations were also transferred to the Ki67 slides and the CCO^−^ patch annotations were duplicated. Duplicates were placed adjacently within WT CCO regions for direct comparison between CCO-proficient and CCO-deficient regions. For PT and CV annotations, 50 μm expansion annotations were generated. Counts for Ki67^+^ cells (DAB + haematoxylin) were performed using the inbuilt ‘Positive cell detection’ tool in QuPath. The PT and CV annotations were also transferred to SOX9+KRT19-stained slides and 50 μm donut-shaped expansions were created. Subsequently, QuPath was trained to distinguish SOX9^+^ KRT19^−^ hepatocytes from SOX9^+^ KRT19^+^ biliary cells and perform counts within the aforementioned donut annotations. Annotation transfers between slides were performed using the script available at https://forum.image.sc/t/interactive-image-alignment/23745/56. Transferred annotations were slightly modified to adjust for the subtle differences in PTs and CVs between serial sections.

### Laser capture microdissection

Laser capture microdissection was performed on 10 μm CCO/SDH-stained cryosections cut onto UV-treated P.A.L.M. membrane slides. Prior to microdissection, slides were completely thawed and air-dried, then tissue was captured in specialised laser-capture tubes (Zeiss, Cat # 415190-9201-000) using a P.A.L.M. laser dissection microscope (Zeiss, Germany).

### Mitochondrial copy number analysis

Tissue and DNA were extracted as per the mitochondrial NGS methodology below, using the same tissue cut sizes and quantities. The Human Mitochondrial DNA Monitoring Primer Set (TaKaRa, Cat # 7246) was used to determine mitochondrial DNA (mtDNA) copy number. Briefly, two mitochondrial (*ND1* and *ND5*) and two genomic genes (*SLCO2B1* and *SERPINA1*) were amplified by qPCR, using 10 μL of template in 50 μL reactions. Amplification utilised Terra qPCR Direct TB Green Premix (Cat. # 638319) and mtDNA abundance was calculated as an average of the two mitochondrial genes relative to the genomic genes using the methodology outlined in the TaKaRa kit.

### Mitochondrial Sanger Sequencing

Single hepatocytes or bulk stroma were laser-capture microdissected and DNA was extracted by incubating the tissue overnight at 65 °C in 14 μL of Arcturus® PicoPure® DNA Extraction buffer (ThermoFisher, Cat # KIT0103). Lysates were then incubated at 95 °C for 10 min to inactivate Proteinase K activity. Amplification of the mitochondrial genome was performed as previously reported (40). Briefly, a nested PCR strategy was used resulting in 36 overlapping amplicons of ~500 bp. This ensured enrichment of the mitochondrial genome at suitable amplicon sizes for Sanger sequencing. PCR products were purified using AMPure XP beads (Beckman Coulter, Cat # A63881) and were Sanger sequenced by Eurofins Genomics (Ebersberg, Germany). All amplicons contained an M13 sequence to facilitate Sanger sequencing. Each sequence file was aligned against the revised Cambridge reference sequence to identify mutations. Sequence visualisations and comparisons were performed using the “A plasmid Editor” available at https://jorgensen.biology.utah.edu/wayned/ape/.

### DNA methylation

DNA Methylation data for the CpG islands in the promoter region of Cardiac-specific homeobox (*CSX*) were generated as described previously (41). Use of a non-expressed gene promotor avoids the potential for expression-related influence on methylation status. Briefly, DNA was extracted and bisulphite-treated, then a 2-stage nested PCR protocol was performed to amplify CpG islands in *CSX*. Successfully amplified PCR products were then cloned to obtained single strand resolution and then Sanger-sequenced.

### Mitochondrial NGS

For each cut site along the portal-central axis of CCO- patches, circles of ~6,500 μm^2^ were laser-capture microdissected. Care was taken to avoid extracting rare intervening CCO+ cells. For each cut position, tissue was captured from ten, 10 μm thick CCO/SDH-stained sections and DNA was extracted using the QIAamp DNA Micro Kit (Qiagen, Cat # 56304), using 2 μL of carrier RNA to enhance DNA yield. Each sample was eluted in 40 μL of warmed H_2_O.

The whole mitochondrial genome was amplified in duplicate, as two overlapping fragments using high-fidelity Takara LA Taq (Takara, Cat # RR002M). Primer details can be found in **Supplementary Table 2**. PCR reactions were performed in 50 μL reaction volumes containing 1X PCR Buffer, 0.4 μM primers, 0.4 mM dNTPs, 1.75 U of TaKaRa LA Taq and 8 μL of template DNA. Reaction conditions were: 94°C for 5 min, then 30-35 cycles of 98°C for 15 sec, 68°C for 10 sec, a slow ramp from 68°C to 60°C at 0.2°C per sec, followed by 60°C for 15 sec, then a 68 °C extension for 12 min. Following cycling, a terminal elongation was performed at 72°C for 10 min. When amplicon abundance was too low for library preparation, samples underwent an additional round of 10-12 PCR cycles using the same conditions. Amplicon quantitation was conducted using Genomic DNA ScreenTapes and reagents (Agilent, Cat # 5067-5365 and 5067-5366) on a 4200 TapeStation System (Agilent, USA)

Following amplification of the mitochondrial genome, libraries were prepared using 1 ng of PCR product, according to the Nextera XT DNA Library Preparation Kit. (Illumina, San Diego CA). Libraries were quantified individually using D5000 ScreenTapes and reagents (Agilent, Cat # 5067-5588 and 5067-5589) on a 4200 TapeStation System (Agilent, USA), were pooled at 4 nM, then sequenced on the Illumina MiSeq v2 and NextSeqTM 500 mid output platforms, both with 300 cycles (150 nt paired-end).

### NGS data processing

Data quality was assessed by FastQC and the mean read depth was calculated for each sample. Samples with low mean read depths were re-sequenced. Reads were aligned to the revised Cambridge Reference Genome (GenBank accession number NC_012920.1) using the BWA-MEM algorithm (42) with a default gap open penalty and gap extension penalty of 6 and 1 respectively. For compatibility with Picard’s MarkDuplicates tool (43), shorter split hits were marked as secondary during read alignment. Aligned reads were then sorted by coordinate using Picard’s SortSam tool, and duplicates were marked and removed with Picard’s MarkDuplicates tool. Single-nucleotide variants (SNV) were identified using the deepSNV R package (44). Bases with Phred quality score of less than 20 (corresponding to an error probability of P = 1% in the original base call) were not considered in SNV-calling. A Benjamini-Hochberg correction was applied for multiple testing, and SNVs with a p-value of less than 0.05 were considered statistically significant.

DeepSNV variants with < 10 supporting reads in the test sample were filtered. Germ-line mutations, defined as those present in both test duplicate samples and in the control at a frequency of > 1%, were removed. Variants not called independently in both technical replicates were discarded. Samples were then grouped according to their patches and portal-central axes. Public variants were defined as those present in all samples along the portal-central axis of a CCO^−^ patch. Conversely, private variants were defined as those which were called in a subset of samples along the portal-central axis.

### Mathematical Modelling and Bayesian Inference to integrate model and data

We performed stochastic simulations of mtDNA mutation accumulation in a two dimensional space using a lattice model (45). In each simulation, cells acquire mtDNA mutations, divide and die. In addition, mutations can occur continuously without cell division as a result of normal cellular mitochondrial replication. Each newly arising mutation is independent and assigned a unique mutation ID. Since our NGS data is not at the single cell level, and thus cannot detect heteroplasmy in individual cells, we only simulate homoplasmic mtDNA mutations in each cell, and do not explicitly model individual mitochondria. Consequently, we assume mitochondria are segregated in equal balance during cell division, and that daughter cells inherit all mutations carried by the mother cell. Model parameters are listed in **Supplementary Table 3**.

To test whether periportal progenitor niches exist, we assign different properties to cells occupying different spatial positions in our lattice model. Hepatocyte-like (non-PT) cells constitute the majority of the system and occupy points on the lattice. Periportal (PT) cells are modelled as a well-mixed cell pool of size N_PT_ in the centre of the lattice. When PT cells divide, their progeny enter the lattice via a cell in the lattice centre and become non-PT cells. Both PT cells and non-PT cells can divide. We assume three alternative scenarios, where PT cells divide slower, faster and at the same rate compared to non-PT cells. The rates of PT and non-PT cell division are related by r^(PT)^ = βr^(non−PT)^, where β≥0 is a model parameter. When β=1, both PT and non-PT cells divide at the same rate and there are no special niches for PT cells. The regime 0 ≤ β < 1 leads to “non-localised” expansion with the mutation accumulation over space dominated by the non-PT cells. Values of β > 1 lead to “localised” dynamics, where mutation spatial pattern is driven by the expansion of PT cells in the lattice centre, which corresponds to the existence of periportal progenitor niches.

By modelling these alternative hypotheses under a range of β values and simulating the mtDNA mutations in cells located in space, we can predict different possible spatial patterns of mutation accumulation as a consequence of different hypotheses. Finally, we compare our spatial patterns estimated by multiple simulations with our NGS data and use Bayesian statistics to infer the most likely range of β for each sample. Detailed methodology can be found in **Supplementary Methods 1**.

## RESULTS

### Characterisation of CCO-deficient clonal expansions

We previously established the use of cytochrome *c* oxidase (CCO) deficiency as a means to investigate dynamics of human tissue clonal cell populations including the liver (32, 33, 46–50). Here, we have utilised this technique permitting the visualisation of blue, CCO-deficient clonal hepatocyte expansions, henceforth referred to as ‘patches’. Observations from large area samples reveal clones extending from the portal tree (**Fig. 1a**). To characterise the patches in our cohort, we first investigated their abundance with age. Detection of CCO-deficiency by enzyme histochemistry requires mitochondrial homoplasmy, or a high degree of heteroplasmy that takes decades to develop stochastically (36). Indeed, in our cohort, the number of patches observed per mm^2^ increased significantly with patient age **(Fig. 1b)**. Despite CCO-deficiency, blue patches were previously determined to be metabolically indistinguishable from their adjacent, brown, non-clonal, CCO-proficient counterparts (32). In this study, we digitally counted the number of nuclei in blue; CCO^−^, and equally sized neighbouring brown; CCO^+^ areas and found the former had significantly more nuclei/area (**Fig. 1c**). Hepatocytes collected by laser microdissection from throughout the lobule had 2.2-fold greater mitochondrial DNA (mtDNA) copy numbers in blue vs brown regions **(Fig. 1d)**. No mtDNA copy number differences were observed when comparing periportal vs pericentral hepatocytes within brown (**Supplemental data 1a&b (S1a&b)**) or blue (**Fig. S1c&d**) regions. We also evaluated proliferation by Ki67 staining; calculated as a percentage of total cell nuclei rather than by area due to the increased cells numbers in blue patches. As expected in normal liver, overall Ki67 positivity was low. We found no significant difference in the proportion of proliferative cells in CCO-deficient patches vs adjacent CCO proficient regions **(Fig. 1e)**.

**Figure 1.**
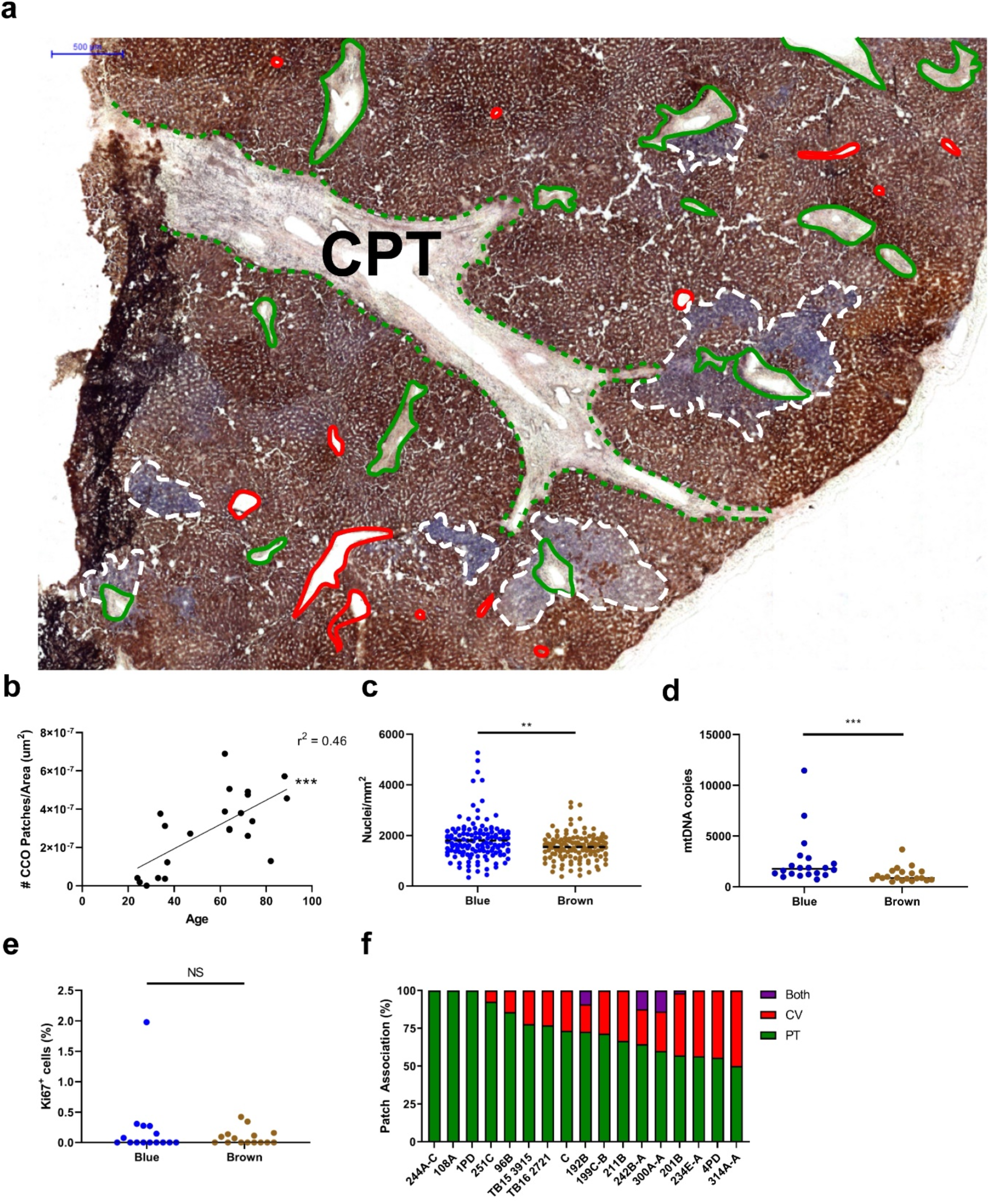
Hepatocyte clonal expansions characteristics. **(a)** A low power image of a CCO/SDH stained liver section with blue CCO-deficient cell ‘patches’ (white border) emanating from portal tracts (green) that branch from the conducting portal tract (CPT; green dotted outline). Central veins are highlighted in red. **(b)** The number of patches normalised by tissue area vs patient age at resection. Data are presented from n=22 patients and 688 patches, \*\*\**P*<0.001; testing by Pearson correlation coefficient. **(c)** The number of nuclei within blue, CCO^−^ and equally sized, brown, CCO^+^ patches from 4 patients, n=130 patches, \*\**P*<0.01. **(d)** The number of mtDNA copies relative to nuclear genomes in CCO-deficient and proficient hepatocytes. Data are from n=20 microdissections each, \*\*\**P<*0.001. **(e)** The percentage of CCO-deficient and proficient cells that are Ki67^+^. Detections within CCO-proficient cells are determined within duplicates generated from CCO-deficient patch annotations that are located adjacent to their CCO-deficient counterparts. Data are from n=13 patients. **(f)** The % of patches in a given patient that were closer PTs, CVs or within 50 um^2^ of both structures; n=17 patients. Statistical tests were performed using Mann-Whitney unless otherwise stated. Bars represent medians.

We next sought to localise the source of these clonal expansions, as stem cells or committed progenitors are thought to be long-lived enough to acquire homoplasmy and thus be cells of origin of CCO-deficient patches (32). This could offer important insights into hepatocyte dynamics in normal human liver. Observationally, CCO^−^ patches appear to emanate from terminal portal tracts (**Fig. 1a**). This propensity for periportal patch localisation was proposed in our previous investigations of histologically normal human liver (32, 33), but never quantified. Here, we have determined the closest structure; portal tract (PT) or central vein (CV), for 463 CCO^−^ patches of ~10 cells or greater in 16 normal human livers **(Fig. 1f)**. In these livers, 74.2% of patches were located closer to PTs than CVs, whilst only 23.6% were closer to CVs. A much smaller proportion (2.2%) were sufficiently large to associate with both structures (**Fig. 1f**). Patch localisation thus favours the presence of a periportal stem/progenitor niche in phenotypically normal human liver.

### Hepatocytes and cholangiocytes share a common somatic ancestor

Should CCO-deficient patches originate periportally, the biliary epithelium or nearby progenitor-like, hybrid hepatocytes (37), may well be a source of new hepatocytes. To investigate this possibility, we utilised CCO-deficiency to locate deficient biliary epithelium with nearby blue hepatocyte patches. **Fig 2a** shows an H&E of morphologically normal liver with distinct portal tracts. CCO-deficiency was identified within the biliary epithelium **(Fig. 2b)** in 3 out of 54 patients. In one of these cases, several CCO^−^ patches adjacent to terminal portal tracts branched from a conducting portal tract containing the CCO^−^ biliary epithelium. Three separate regions of CCO-deficiency (**Fig 2b** labelled area 1; biliary epithelium and areas 2 & 3; hepatocytes) were laser-microdissected and examined for clonality **(Fig. 2c&d)**. A common *m.8251G>A* mutation (in *MT-CO2* coding for subunit COII of cytochrome *c* oxidase) was observed between the CCO-deficient biliary epithelium in LCM area 1 **(Fig 2c&g; ii&iii)** and CCO-deficient hepatocytes within LCM area 3 **(Fig. 2d&h; iv)**. This mutation was not present in the stroma **(Fig. 2e)** from the same patient, nor was it detected in the majority of nearby CCO-proficient hepatocytes tested **(Fig. 2f)**. Two CCO-deficient hepatocytes tested within LCM area 2 were found to be clonal by virtue of a shared *m.1214A>G* mutation, but they did not contain the *m.8251G>A* mutation present in the CCO^−^ duct **(Fig. S2)**. The chances of a common mutation independently occurring in these two cells is extremely low; 2.48×10^9^:1 (33, 51). As such, these data demonstrates a likelihood for the existence of a periportal bipotential common ancestor of biliary epithelial cells and hepatocytes, and to our knowledge, is the first such report in normal human liver.

**Figure 2.**
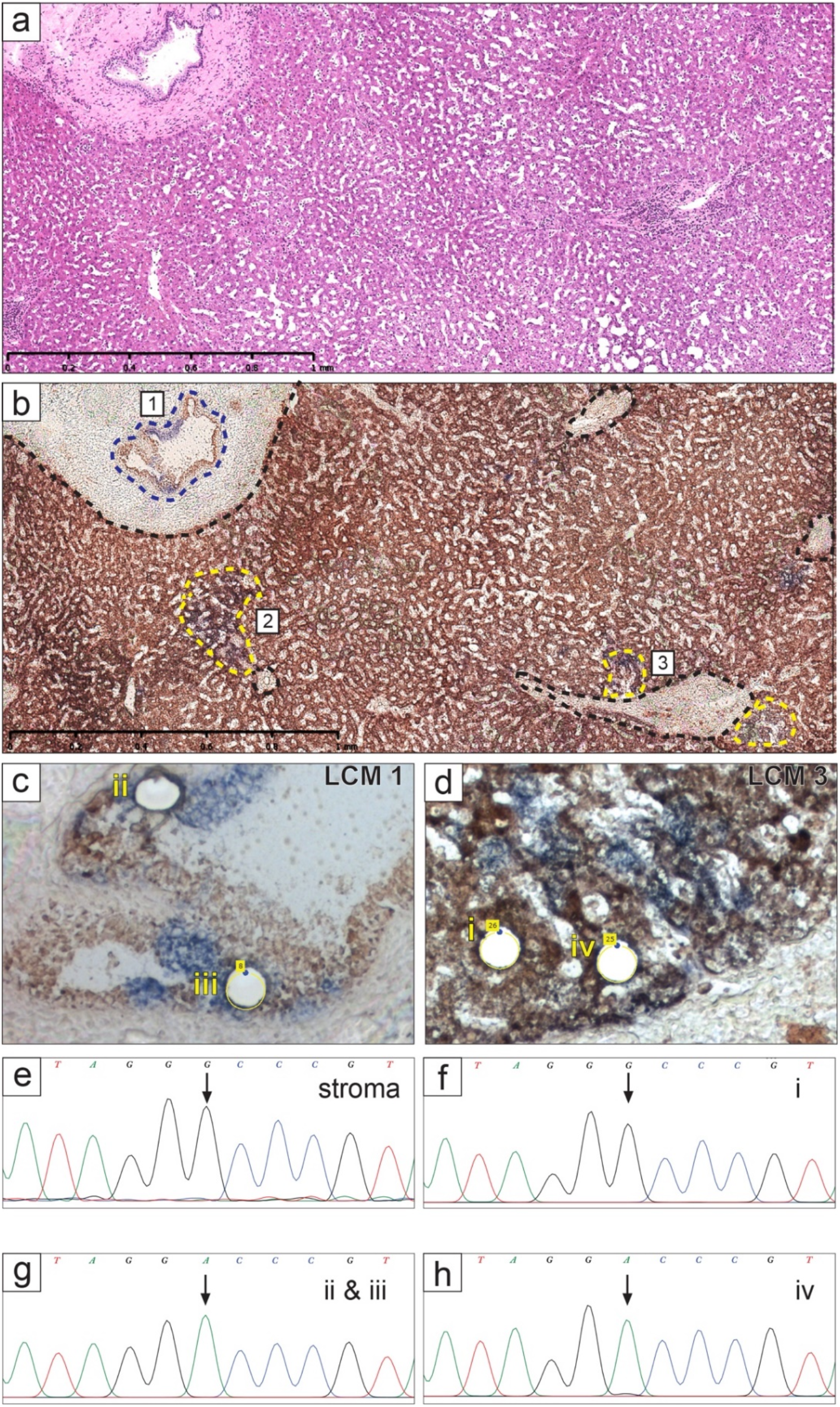
An ancestral relationship between biliary epithelium and hepatocytes. **(a)** An H&E **(a)** and **(b)** a serial CCO/SDH-stained section highlighting three (labelled areas 1-3) microdissected regions. Portal tracts as well as CCO-deficient hepatocyte patches and biliary epithelium are outlined in black, yellow and blue respectively. **(c)** High power image of LCM area 1 from **(b)** displaying two laser microdissections of CCO-deficient biliary epithelium (ii&iii). **(d)** A high-power image of a laser-microdissected CCO-normal (i) and CCO-deficient (iv) hepatocyte from LCM area 3. **(e-h)** Electropherograms from Sanger sequencing of distal stroma, a CCO-proficient hepatocyte, CCO^−^ biliary epithelium and CCO^−^ hepatocyte. Arrows designate the location of the *m.8251G>A* mutant.

### SOX9+ and Ki67+ hepatocytes are present in normal human liver and are predominantly periportal

Normal liver is largely quiescent, however should hepatocyte turnover in this tissue be actively fed from periportal cells, this region may be more proliferative than at the CV. To investigate this, we annotated all PTs and CVs within normal livers from 13 patients using the pathology software QuPath, and we detected Ki67^+^ cells within 50 μm expansions from the boundaries of these structures **(Fig. 3a&b)**. We observed no significant increase in the proportion of Ki67^+^ cells at the PT vs CV **(Fig. 3c)**. We also investigated the possibility of a SOX9^+^ progenitor as a source of hepatocyte clonal expansion (37). As SOX9 will also stain ducal cells, we stained slides by dual IHC for SOX9 and KRT19 **(Fig. 3d&e)** in order to exclude SOX9^+^ these. QuPath was then trained to differentiate between SOX9^+^ KRT19^+^ ductal cells and SOX9^+^ KRT19^−^ hepatocytes within 50 μm expansions surrounding CV and PT. SOX9^+^ hepatocytes were detected in significantly greater abundance within 50 μm of PTs relative to CVs; a mean of 14.85 vs 5.89 cells/mm^2^ **(Fig. 3f)**. This is the first report of a periportal preponderance of SOX9^+^ hepatocytes in normal human liver. Collectively, these results support a progenitor niche at this anatomic location.

**Figure 3.**
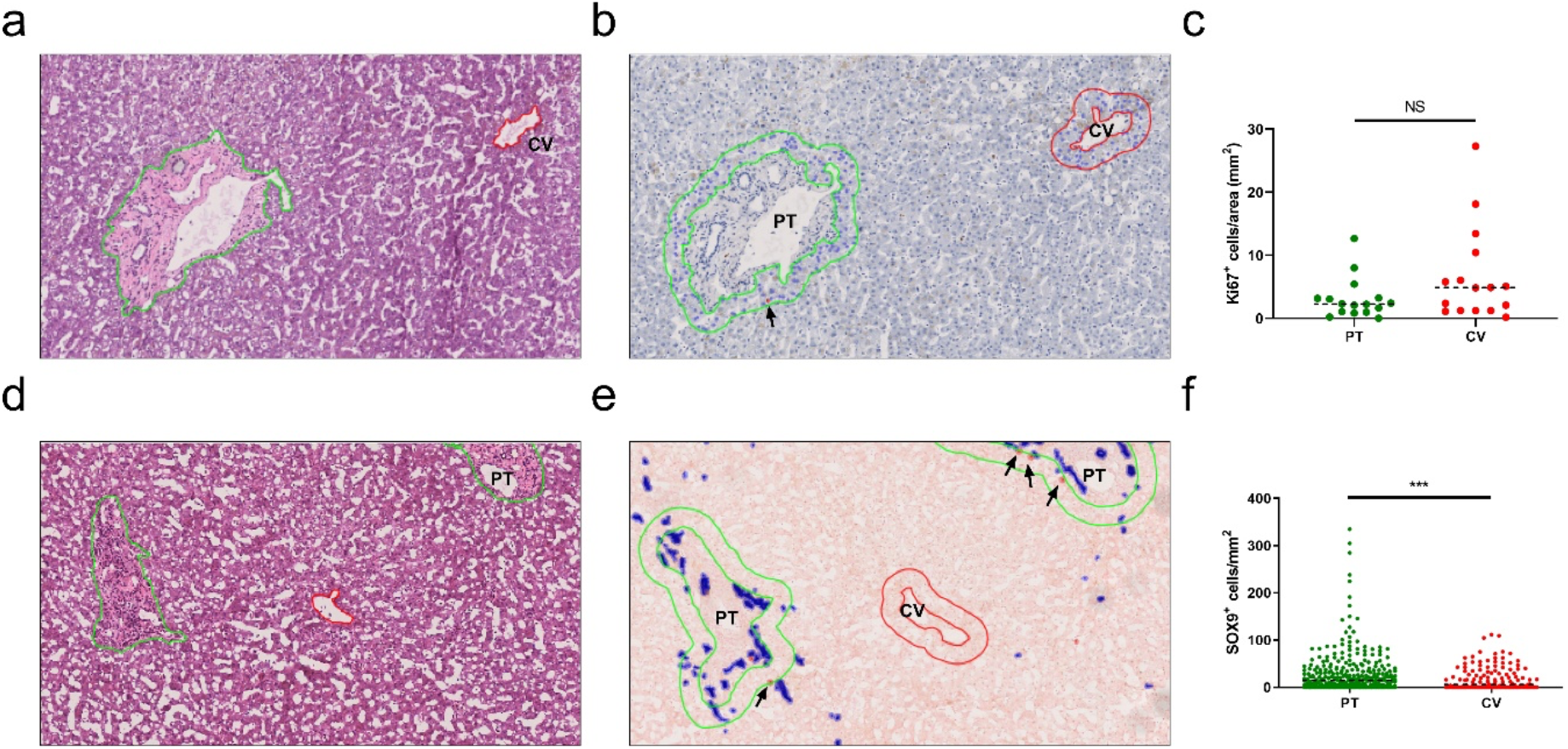
SOX9^+^ hepatocytes are present in normal human liver and are predominantly periportal. **(a&d)** H&Es with **(b)** accompanying Ki67 IHC and **(e)** SOX9/KRT19 dual IHC. **(b&e)** Portal tracts (PT; green) and central veins (CV; red) were annotated then expanded by 50 μm to permit detection within an area surrounding these structures. Red cells denote detections of Ki67^+^ and KRT19^−^ SOX9^+^ cells and arrows indicate detections within the 50 μm expansions that counted towards the analysis. Blue detections in **(e)** are KRT19^+^ SOX9^+^ biliary epithelium. **(c)** Density of Ki67^+^ cells within 50 μm PT and CV expansions from n=16 patients; NS=not significant by Mann-Whitney test. **(f)** Density of SOX9^+^ hepatocytes within 50 μm expansions from PTs and CV boundaries within 6 patients for n=602 and 430 PTs and CVs respectively. Bars represent means and *P*\*\*\*<0.001; Mann-Whitney.

### Methylation diversity supports a periportal stem cell niche

Given our evidence for a periportal progenitor niche, we next sought to investigate the expansion dynamics of clonal patches in normal human liver. For this, we used methylation diversity as a dynamic clonal marker – one that changes with time and size of clonal expansion (41, 52). Methylation patterns record cell ancestry as they are inherited with high fidelity by daughter cells. The greater the methylation diversity between cells, the more distant the ancestral relationship (52). We used two markers of diversity: intra-patch distance (calculated as the minimum number of sequence changes required for each sample to have the same sequence) and the number unique methylation sequences (tags). First, we measured methylation diversity within large periportal and pericentral regions without regard to clonality and found no difference in the epigenetic distance between groups **(Fig. S3a-c)**. Next, CCO-deficient periportal, midzonal and pericentral cuts were analysed from each patch and no differences were observed **(Fig. S3d-f)**. If the streaming liver hypothesis (19) is correct, and hepatocytes migrate along the portal-central axis, the ancestral and epigenetic distance should increase with anatomical distance from the stem cell niche. Cuts were then made within the patches, at varying distances from the PT along the portal-central axis **(Fig. S3g-i)**, but again no correlation was observed between epigenetic and anatomical distance. From these data, it appeared the epigenetic diversity measurable using this technique was maximised within each tissue extraction.

Smaller extractions, remaining close to the putative stem cell niche are more likely to remain within the limits of this diversity measure. To this end, we investigated methylation diversity in patches of increasing size that abutted the portal tract. Measuring diversity is heavily influenced by the number of cells analysed and to control for this we micro-dissected the same sized area from each clone. Larger clones had multiple dissections; each of the same size, but each was analysed independently. Extractions were made within small, medium and large CCO^−^ patches of sizes, directly abutting PTs **(Fig. 4a&b)** and CVs. Methylation tag diversity increased with patch-abuttal size at the PT **(Fig. 4c)**. A positive correlation with a slope significantly deviated from 0 was observed between epigenetic distance and the length of the patch abutting the PT **(Fig. 4d)**. No such correlation was observed for patches abutting the CV **(Fig. 4e)**. These data demonstrates an ancestral relationship between hepatocytes at the PT but not the CV that dissipates with the expansion of the patch.

**Figure 4.**
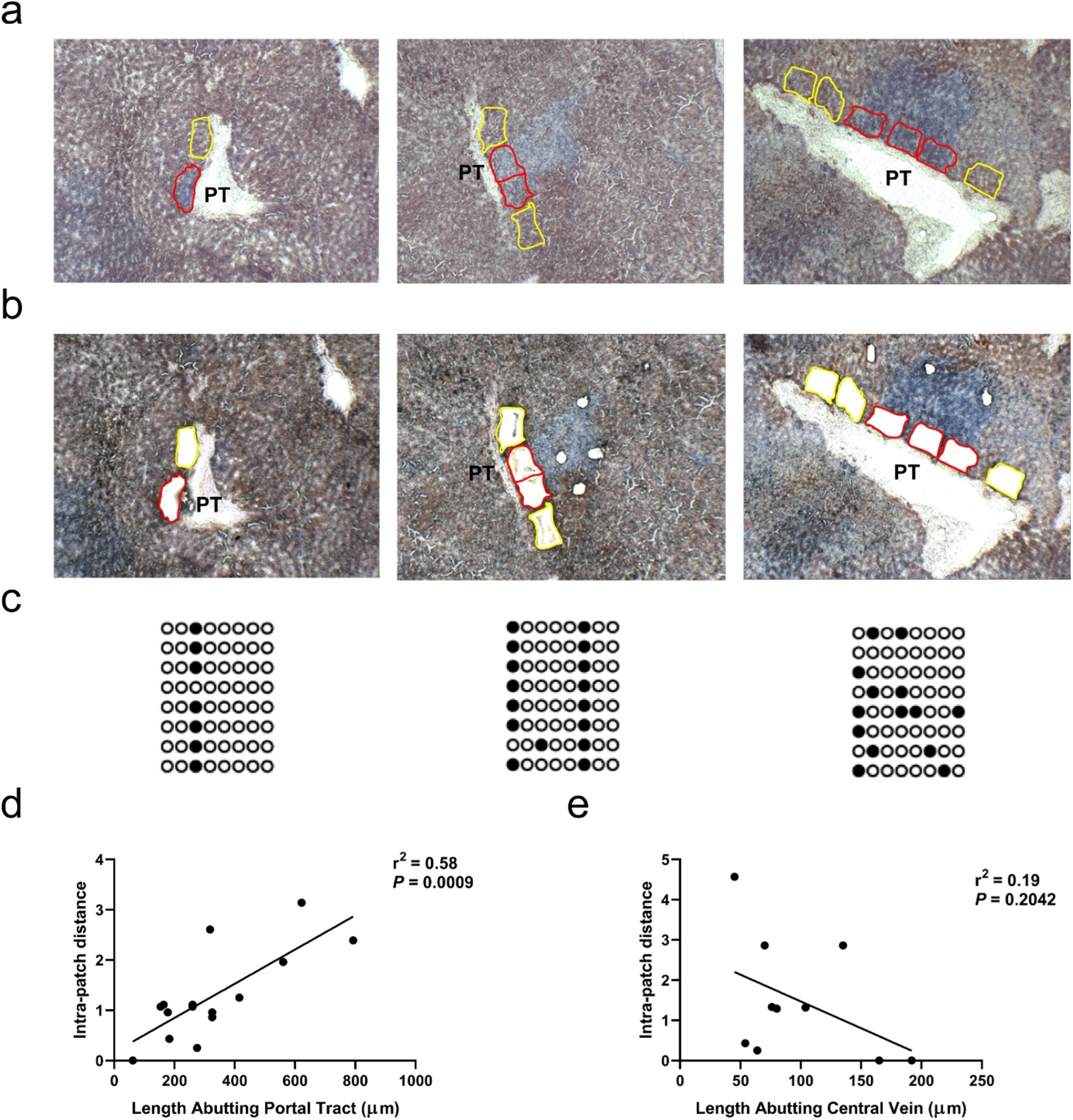
Methylation diversity demonstrates a periportal hepatocyte origin**. (a&b)** From left to right; representative cytochrome *c* oxidase-deficient (blue) patches of increasing abuttal length, before **(a)** and after **(b)** laser-capture microdissection. Red and yellow outlines highlight the equally-sized CCO-deficient and CCO-proficient regions captured. **(c)** Lollipop diagrams for the three patch sizes abutting portal tracts. Each column represents a CpG site and each row represents a sequenced cell. Circles are solid when a CpG site is methylated. **(d&e)** Methylation diversity vs patch abuttal length correlations obtained from n=15 portal tracts and 10 central veins. Statistics calculated using Pearson correlation test.

We hypothesise there are no differences in expansion dynamics between CCO-deficient and proficient patches. Indeed, we found no significant difference in the % methylation of our CpG sites between CCO-deficient and CCO-proficient cells **(Fig. S3j)** indicating patch dynamics are similar. Given the relationship determined within known CCO^−^ clonal boundaries, we also measured methylation diversity in size and spatial location-matched CCO-proficient PT and CV abutting areas. Whilst a weak correlation of intra-patch distance of patches of increasing size was observed in CCO^+^ areas, this was weaker than in CCO^−^ patches, and the slopes were not significantly deviated from zero at the PT or CV **(Fig. S3k)**. This demonstrates the benefit of examination within known clonal boundaries, which are otherwise difficult to determine in normal liver. Similar overall results were achieved using another measure of diversity; the simple number of unique methylation patterns or ‘tags’ observed within a given extraction **(Fig. S3c, f, i, l & m)**.

### Clonal hepatocytes display large numbers of unique mitochondrial DNA variants across PT-CV axes

Our data thus far indicated the presence of a periportal progenitor niche being the origin of CCO-deficient patches, however the spatio-temporal dynamics of these expansions could not be determined with the limited resolution of our methylation assay. To address this, we performed deep NGS of mtDNA genomes, a technique with far greater resolution, enabling analysis across the whole patch. In a similar fashion to methylation diversity, mtDNA mutations can be exploited to determine the relatedness of cells across a patch. As mitochondria will continue to replicate in the absence of cell division, mutations can accumulate even in largely proliferatively quiescent tissues such as the liver.

We would hypothesise that if hepatocytes were continuously streaming from their cell of origin, we would be able to detect a distinct stepwise pattern of inheritance of mtDNA variants as we sample across each patch. Older cells, hypothetically at the CV, would have had more time to develop independent mitochondrial variants and would be more dissimilar to their neighbours compared with cells at the PT origin. Alternatively, should patches grow in a discontinuous fashion, with intervening quiescence, we would observe a large degree of independent variants across the patch.

To ensure no large mtDNA copy number differences exist between periportal and pericentral hepatocytes, mitochondrial copy number was determined at these locations in CCO^−^ hepatocytes **(Fig. S1c&d)**. No significant differences were observed. Similarly, no differences were observed between CCO^+^ PT and CV hepatocytes **(Fig. S1a&b)**.

Four to five ‘cuts’ of size-matched hepatocytes were laser microdissected along the PT-CV axis in CCO-deficient patches that bridged PTs and CVs **(Fig. 5a)** After DNA extraction, mtDNA genomes were sequenced by NGS and the variants within each cut were called. In total, 68 cuts across 14 PT-CV axes were extracted from 10 patches across five patient samples. The loci of all PT and CV variants were also plotted for each patch and no discernible differences were observed **(Fig. 5b).** The mutation types for cuts adjacent to PTs and CVs were compared to investigate whether differences between these locations could be observed **(Fig. 5c)**. Overall, the base substitutions were very similar, with the T→C substitution predominant in cuts from both locations. This substitution has been associated with aging (53), a phenomenon coinciding with the development of the large patches used for this analysis **(Fig. 1a)**. Accurate mutational signatures could not be generated due to a low total number of somatic variants detected (771 SNVs) across our samples.

**Figure 5.**
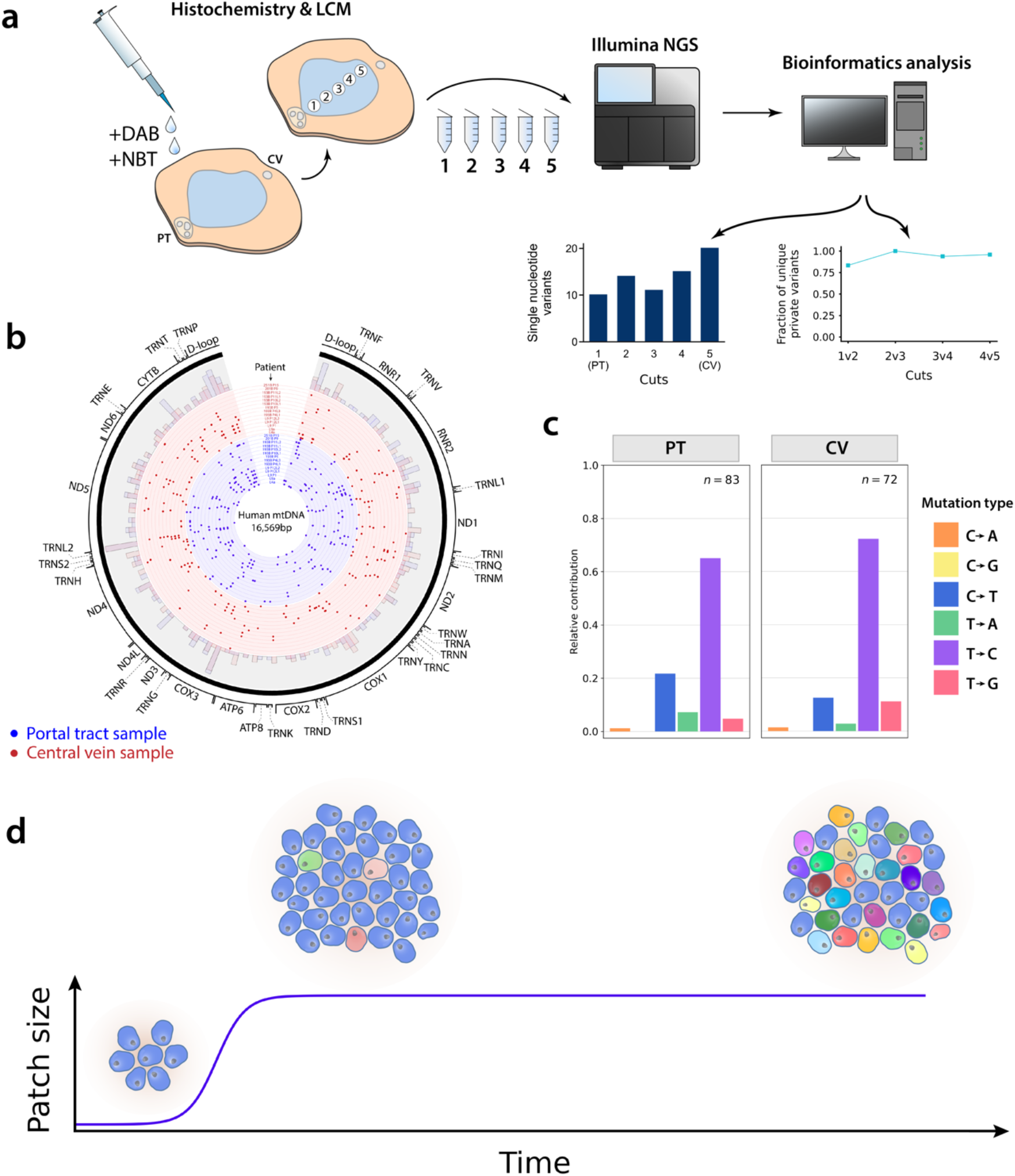
Next-generation sequencing of CCO-deficient hepatocytes. **(a)** Typical NGS workflow. A CCO/SDH stained section is laser microdissected across the portal tract (PT) to central vein (CV) axis. Samples along the axis are sequenced and analysed individually. **(b)** Distribution of single nucleotide variants across the mt-genome from 68 laser-microdissected cuts across 14 PT-CV axes in 10 patches from 5 livers. The innermost section (blue data points) represents data for portal tract abutting samples and outer rings (red data points) represent central vein abutting samples. The combined SNV data for all portal tract and central vein abutting samples is shown in a histogram (100bp bin width). **(c)** Single base substitutions aggregated by type observed in all PT and CV cuts. **(d)** A schematic for the proposed model of clonal expansion of hepatocytes in normal human liver. A patch of clonal cells expands, then quiesces for extended time periods. With time, each cell acquires unique mtDNA variants (represented by cells with different colours) which develop independently of cell division.

Most PT-CV axes contained a large percentage of unique private variants; (not shared by neighbouring pairs of cuts) **(Fig. 5a)**. By virtue of their shared CCO-deficiency, we know that all cells within a given patch share a common ancestor; yet these data indicate cells in each cut are largely genetically distant from their physical neighbours. We propose that this pattern is suggestive of punctuated expansion of clonal patches, with extensive periods of quiescence. This would allow sufficient time for the cells in each cut to develop variants that are unique when compared to their neighbours **(Fig. 5d)**.

### Spatial modelling supports neo-hepatocyte generation from a periportal niche

We built a spatial simulation model of liver tissue dynamics to explore scenarios that further test our hypothesis of a periportal niche. We compare all possible mtDNA spatial patterns predicted under different model assumptions with the patterns observed in NGS data **(Fig. 5a)**. The model explores two possible modes of clonal expansion: homeostatic cell turnover dominated by proliferation of randomly distributed ‘non-PT’ parenchymal cells, or alternatively, faster replication by a small number of stationary ‘PT cell’ progenitors at the centre of the model **(Fig. 6a)**. These two regimes are described by our model parameter ‘β’, with the value of β<1 and β>1 respectively. Rounds of homeostasis take place characterised by the death of randomly located non-PT cells with subsequent replacement and mitochondrial mutation **(Fig. 6b)**. Cell death leaves open space, allowing for occupation during cell replacement. An adapted cell-pushing algorithm (45) is applied whereby the daughters of dividing cells ultimately occupy a neighbouring space, opened by displacing neighbouring cells into empty spaces made available by prior cell death **(Fig. 6c)**. After each round of homeostasis, non-PT cells in discrete zones along a random radial axis are sampled to record their mutational burden **(Fig. 6d)**. As the absolute size of mutational burden measured depends on the number of cells sampled (N_s_) per zone, we used N_s_ = 60 in accordance with the average number of cells in each cut of our NGS experiment. As described in the **Methods**, our model also captures the phenomenon of mitochondrial mutations without cell division. We used a literature based mitochondrial mutation rate of 0.1 per cell division (54, 55). Extensive details are in **Methods** and **Supplementary Methods 1**.

**Figure 6.**
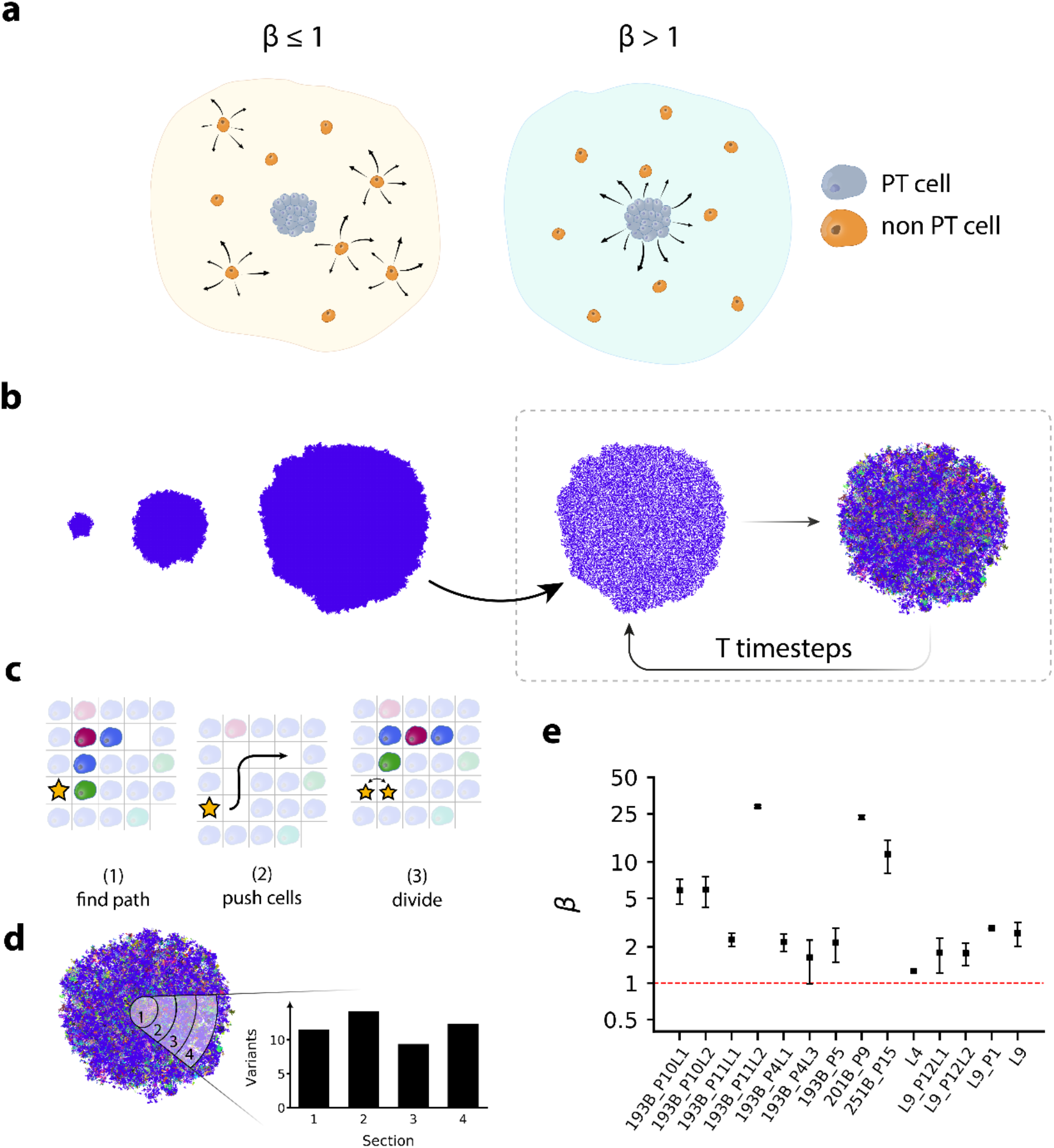
Spatial modelling of homeostatic liver dynamics. **(a)** A low (≤1) β regime has non-localised dynamics in which divisions of non-PT cells occur at a faster rate and dominate the dynamics. A β > 1 leads to localised dynamics, in which divisions are dominated by a small number of PT cells in the centre of the system. **(b)** Simulation phase 1; a single cell seeded followed by simulated cell divisions. The pool of PT cells is at the system’s centre. Cells are coloured by genotype, with all cells in the initial system beginning with zero mutations. For ‘T’ timesteps, 50% of cells selected at random, are killed, and the remaining cells divide and repopulate the system. Cells acquire new mutations resulting from these divisions. **(c)** Our cell-cell pushing algorithm; a dividing cell (star) finds a path to a nearby empty lattice point. It pushes cells along this path to create an empty space next to itself, then divides, with one of its progeny occupying the newly created empty lattice point. **(d)** Cells are sampled in “sections” according to radial distance from the centre of the system. Sections are located along a randomly orientated radial axis passing through the system’s centre. The central section is denoted “section 1”, with sections 2–4 lying at increasing radial distances from the central zone. PT cells reside in the central section, as a well-mixed population. The number of unique variants, occurring at a frequency above the detection threshold, is plotted for each section. **(e)** An Approximate Bayesian Computation rejection sampling algorithm of β was run for each portal-central axis assayed by mitochondrial NGS and the geometric median point estimates of β are displayed. Patient codes and sample names, respectively, are shown in the legend. Errors are S.E.M.

Each simulation was run over 100 rounds of homeostasis under various combinations of β and number of PT cells (N_PT_). To compare our simulated zonal mtDNA mutation patterns with those along each PT-CV axis obtained in our experimental NGS data **(Fig. 5a&b)**, Approximate Bayesian Computation (ABC) rejection sampling was implemented. This was applied to estimate the joint posterior distribution of the β and N_PT_ parameters that best simulate our experimental NGS data. Approximately 2,000 β and N_PT_ parameter pairs were sampled from the posterior distribution for each experimental sample, and geometric medians were used as a point estimator for the best-suited combination of parameters **(Fig. S4-17)**. Geometric medians were then collectively plotted to visualise any unifying patterns in our data. Overall, all experimental samples are best fit by β>1, which signifies the presence of a small number of static ‘PT’ progenitors that replicate faster than the randomly distributed parenchymal, ‘non-PT’ cells **(Fig. 6e)**. As we observe a low sensitivity to the N_PT_ parameter in our model, we did not infer the number of PT given our current dataset. Collectively, this modelling supports our hypothesis for the generation of CCO^−^ patches from a confined niche and not solely from randomly-distributed hepatocytes.

### Clonal expansions expand slowly and become quiescent

We tested our expansion dynamics hypotheses by again simulating the distribution of mtDNA variants, but in this instance, across a simple one dimensional, in silico PT-CV axis (**Supplementary Methods 2**). The simulation takes place over two phases; first, a clonal expansion occurs either slowly or rapidly to populate the axis. As was determined in our prior model, generation of new hepatocytes by the PT-cell positioned at the simulated portal tract occurs more rapidly than by randomly distributed hepatocytes along the axis. At the conclusion of clonal expansion, a second phase consisting of either quiescence or a streaming liver occurs **(Fig. 7a&b)**. As the period between clonal expansion and observation of the patch within the laboratory is unknown, the second phase is simulated across various time periods (0 d, 1 mo, 5 yr and 20 yr). During simulated quiescence all cells stop dividing, however mtDNA mutation persists as they are capable of replication in the absence of cell division. Under streaming dynamics, PT-cell replication persists and as new hepatocytes are generated, those at the simulated central vein are pushed out and lost from the system, akin to an active conveyor belt of hepatocytes. Extensive methodological details and parameters can be located in **Supplementary Methods 2**.

**Figure 7:**
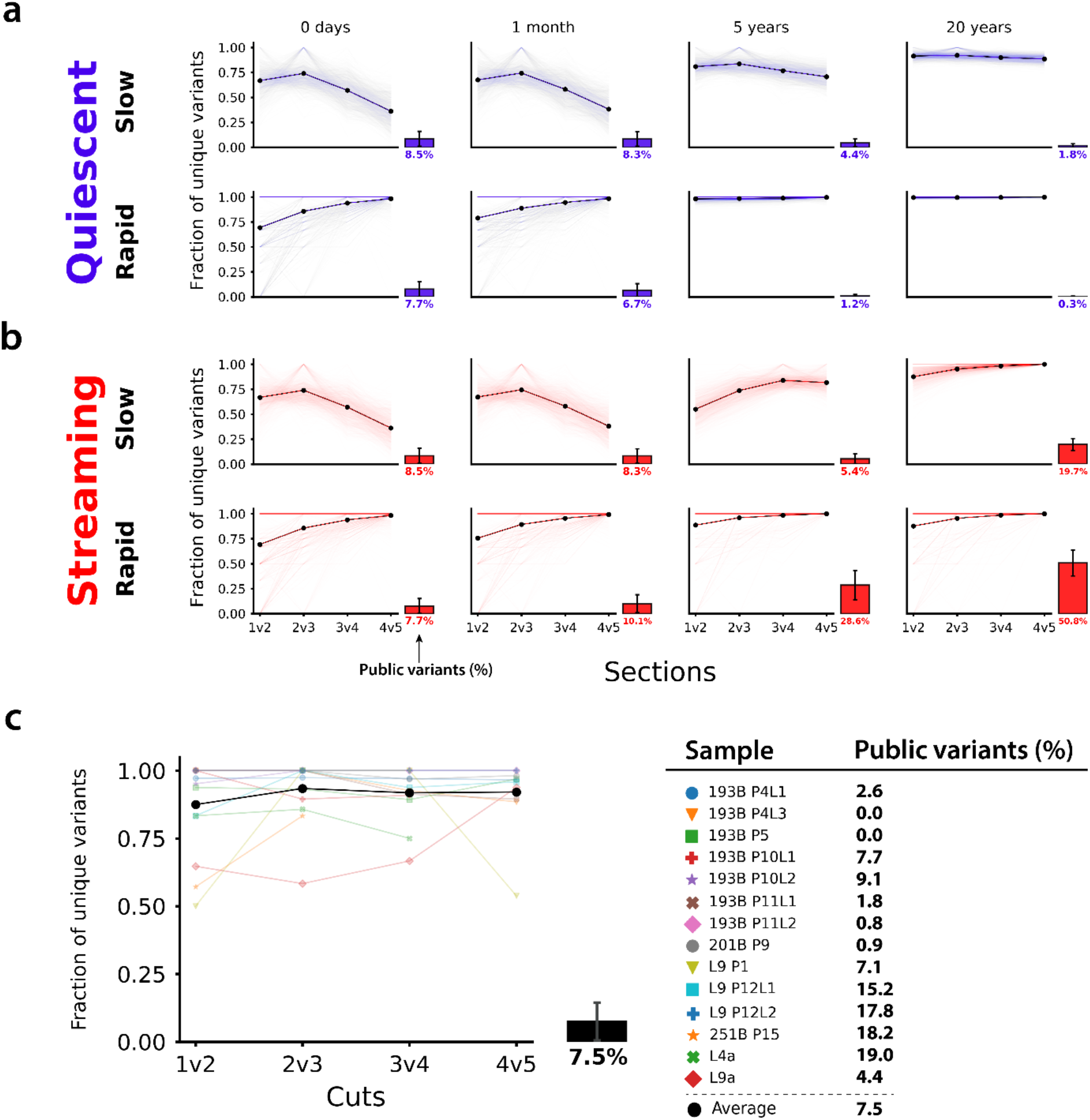
Simulated public and unique private variant dynamics for **(a)** quiescent and **(b)** streaming liver dynamics assuming either slow or rapid initial patch growth. Unique private variants **(y-axes)** are 1) non-public and 2) not shared between adjacent pairs of simulated sections or actual patch cuts **(x-axes). (a&b)** Each panel displays a result based on simulations of 1000 realisations of a 1-dimensional system akin to portal-central axes. Clonal hepatocytes are first generated at either a slow or rapid rate, followed by 0 days, 1 month, 5 and 20 years of quiescent or streaming dynamics. Transparent curves correspond to data for individual realisations of the stochastic system, and the solid curve represents the mean value across all simulations. The proportion of public variants; those shared across all cuts is displayed as a bar alongside each panel (mean ± SD). **(c)** Data for non-simulated experimental samples and their average.

At the conclusion of both phases, the axis is divided into 5 sections of equal cell number, mimicking the patch cuts in the experimental data **(Fig. 5a)**. Unique private variants are detected within each section and compared in neighbouring pairs as in our experimental data. In our simulations, regardless of phase 2 dynamics, the proportion of unique variants in each pair of sections approaches 1 (100%) as the time since phase 1 (clonal expansion) increases **(Fig. 7a&b)**. Interestingly, quiescent dynamics **(Fig. 7a)** serve to decrease the proportion of public variants (those shared across all sections) over time whilst a conveyor-like streaming dynamic **(Fig. 7b)**, achieves the opposite in this respect. In all instances, an initial rapid expansion appears to accelerate the trend formed under a slow expansion system. Whilst there is some variability in the unique variant plots obtained from our NGS data **(Fig. 7c)**, most approach 100% unique variants across the patch. This is most closely represented by longer phase 2 (5 and 20 yr) simulations. Additionally, the experimental data has public variant proportions largely <10%, which is most consistent with quiescent phase 2 dynamics. To determine the simulation that most closely resembled our average NGS data (Fig. 7c; Black line & boxplot) we found the simulated scenario which minimised the Euclidean distance to the average NGS data with respect to the unique private variants curve and public variant proportion. A slow expansion with 20 years quiescence was determined to be the best fit for both the unique private variants and public variants data. As such, for the average patch, this modelling lends support to a slow/punctuated clonal expansion with long periods of quiescence.

## DISCUSSION

Most of what is known regarding hepatocyte clonal dynamics in liver is understood from the response to injury models in rodents. Recent lineage tracing techniques have greatly enhanced our knowledge in this regard, however, results in normal liver remain contentious and contradictory. Understandably, there are far fewer options for lineage tracing in humans, resulting in an even wider knowledge gap. Large clones dominate within diseased human liver but are much smaller in normal human liver (31). Understanding the expansion dynamics in normal human liver thus benefits from marking clonal patches to avoid cross-clonal interrogation. We address this by leveraging cytochrome CCO deficiency to delineate these smaller clones, and in doing so, we can accurately investigate hepatocyte clonal dynamics. We measure the clonal expansion dynamics by methylation diversity and mtDNA variants through next-generation sequencing of hepatocyte patches sampled spatially. We build mathematical spatial models to simulate tissue growth with or without faster proliferation in a portal-associated niche. Finally, we use Bayesian inference to identify the more likely assumption by comparing our computational simulations and NGS data.

In our study, CCO^−^ patch and SOX9^+^ hepatocyte proximity to the PT, methylation diversity analysis and spatial modelling all support the existence of a periportal progenitor niche for the clonal expansions of hepatocytes. First, quantification of the proximity of clonal patches to CVs or PTs shows that significantly more are periportally located, suggesting this as their origin. As CCO deficiency likely initially develops in long-lived stem/progenitors (33), this localisation would designate a periportal progenitor niche. In some cases, patches may associate with a portal tract that is out of our plane of sectioning. These results alone cannot exclude that a small number of clones may arise from the central vein. However, as additional support in favour of a periportal progenitor niche, we demonstrate a clonal, common somatic ancestor of the biliary epithelium and hepatocytes. This finding raises the likelihood of a periportal bipotential progenitor as a source of clonal hepatocyte expansions. We also identify the presence of putative SOX9^+^ KRT19^−^ progenitor-like hepatocytes (37, 56) in greater abundance immediately surrounding portal tracts. Using methylation as a marker of diversity, we were able to establish a relatedness of hepatocytes at the PT that was absent at the CV, indicating the PT as a site of origin for clonal expansions. Furthermore, the pattern of mtDNA SNVs detected across PT-CV axes within clonal patches is best recapitulated by our computational modelling scenario that supports a disproportionately large portal progenitor contribution to expansion. Collectively our data supports a periportal origin for clonal expansions.

The hybrid hepatocytes (HybHPs) described by Font-Burgada et al. (37) are a good candidate progenitor for these clonal expansions. First, HybHPs represent a pool of rare periportal cells that remain as such until faced with pericentral damage. Second, as the name suggests, HybHPs express a mix of biliary (such as SOX9) and hepatocyte markers. Our data both demonstrated the existence of a somatic bipotential progenitor of hepatocytes and cholangiocytes, and the presence of periportal SOX9^+^ KRT19^−^ hepatocytes.

In addition to determining the origins of hepatocyte clonal expansions, our experiments offer insights on the temporal nature of clonal expansions. Changes in methylation are realised only through cell replication over time (41). This technique, in theory, can then also inform us on expansion dynamics. Should a patch expand, and stop, there is insufficient replication for the methylation pattern to appreciably alter, thus the diversity of patterns observed across the patch should remain low. Subsequent quiescence, as is generally observed in normal liver, would ensure that the methylation diversity remains largely unaltered. A continuous streaming scenario would involve a greater amount of replication, in theory allowing differences to be observed between the cell origin and terminus. Unfortunately, our methodology did not have the resolution to distinguish these expansion dynamics. This could be improved by methylation systems that allow a greater number of CpG sites to be sampled within a given target promoter. Regardless, our methylation technique was beneficial for its support of the periportal clonal origins as mentioned.

By contrast, mitochondria will replicate in the absence of cell division and develop SNV diversity over time, even within quiescent cells. Patch expansion followed by lengthy quiescence allows even neighbouring clonal cells to largely individualise their mtDNA variant profiles, greatly reducing the proportion of public variants; those present across the whole PT-CV axis. Alternatively, a conveyor-like streaming liver model (19, 37) serves to greatly increase the proportion of public variants over time. Both these hypotheses are confirmed by our simulations. Within our experimental data, the average patch has a low level of public variants with largely individualised variant pools, which is best matched by our long-term (~20 years) quiescence and slow patch expansion simulations. The slow patch expansion scenario translates to a periportal cell replication rate approximately 3 times greater than that of randomly distributed hepatocytes in the 1-dimensional model: a value broadly consistent with the Bayesian estimates from our 2-dimensional modelling. An untested but plausible scenario that may still explain our data would be punctuated streaming with long periods of quiescence. Furthermore, limiting the simulations to 1 spatial dimension will likely accentuate the effects of streaming dynamics. If hepatocytes do exhibit streaming *in vivo,* they would not always strictly travel along the 1-dimensional PT-CV axis. These factors may explain the variance observed at an individual patch level and the discordance between experimental measurements and the simulated data generated under streaming dynamics.

Questions remain as to the cause of these patch expansions. As CCO deficiency is not expected to cause a selective advantage (36), it is reasonable to assume that these clonal expansions are a liver-wide phenomenon; not restricted to those made visible by CCO/SDH staining. Our methylation data indeed indicates that CCO-proficient and deficient hepatocytes are equally replicative. Additionally, CCO-proficient and deficient hepatocytes are indistinguishable in terms of Albumin, cytochrome P450 1A2 staining and Ki67 (32). Given we are analysing histologically normal tissue, these clonal expansions could indeed be a homeostatic process. Our current understanding of this process is almost exclusively rodent-based and contradictory. There is evidence for homeostatic hepatocyte turnover from specific liver zones (17, 22, 24) as well as from non-zonal, randomly distributed hepatocytes (25, 26). There is, however, a consensus that hepatocytes are relatively quiescent during homeostasis. Our data supports a combination of dominantly periportal patch expansion and long-term quiescence. It remains possible that clonal patches result from repeated bouts of acute injury from environmental or lifestyle exposures/stimuli. Indeed, pericentral injury, often initiates a periportal replicative response (37, 57), however, chronic injury is usually required for new hepatocytes to extend far beyond this niche. The majority of our samples came from resections distal to metastases, thus many patients had received systemic chemotherapy, but other lifestyle data were not collected. Examining clonal patches from younger donors who have yet to experience repetitive hepatotoxic injury could in theory shed further light on this matter, however normal tissue from this age cohort is extremely rare. Additionally, our lineage tracing methodology, combined with our selection criteria of patches bridging the PT and CV, by its nature, predisposes our analyses to older patients who have more abundant patches, but are unlikely to have had a life free from damage. Indeed, the youngest donor used in our NGS analysis was 62 years of age at time of resection.

Hepatocyte clonal dynamics in humans has been vastly under-studied. Knowledge of these dynamics in normal human liver is essential as baseline for comparison to disease responses. Furthermore, identification and characterisation of a progenitor niche could ultimately be therapeutically beneficial. We have demonstrated the presence of a periportal progenitor niche in phenotypically normal human liver and have provided evidence that neo-hepatocyte expansions spread from this niche. Further work is required to understand what initiates these expansions and how their dynamics may be altered.

## Supporting information

Supplemental tables, methods & results

## Acknowledgements

We are grateful for Members of Cancer Tissue Bank (https://www.cancertissuebank.org) operational team (Amina Saad, Thomas Dowe, Ahmet Imrali, Abhirup Banerjee, Claude Chelala, Sameena Iqbal, Dayem Ullah, Rory Smith, Catherine Graham, Sarah Mueller) and Tissue Access Committee (Satyajit Bhattacharya, Chairman) for the valuable support in this project (2015/01/QM/SM/E/FrozenTissue).

