## Supplemental tables, methods & results for "Clonal dynamics of normal hepatocyte expansions in homeostatic human livers and their association with the biliary epithelium"

**Supplementary Table 1 - Patient Details**

| Anonymised code | Age (yrs) | Sex | Reason for resection | Chemotherapy |
|---|---|---|---|---|
| C | | | No data available | No data available |
| C01P0188B KK01 | 21 | F | Mucinous Cystic Neoplasm | None |
| C01P0334B KK11 | 24 | M | Colorectal Liver Metastasis | Chemotherapy received [last dose 18/05/2019] |
| C01P0314A KK06 | 25 | M | Hytadid Cyst | None |
| C01P0104B KK02 | 28 | M | Gastro-intestinal Tumour of the stomach, liver metastasis. Carrier of Familial SDHB mutation | None |
| C01P0211B KK05 | 33 | F | Uveal Melanoma Liver Metastasis | None |
| C01P0192B KK06 | 34 | F | Focal Nodular Hyperplasia | None |
| C01P0244A KK06 | 36 | M | Hepatic Adenoma | None |
| C01P0263A KK06 | 36 | M | Colorectal Liver Metastasis | XELOX [started 11/05/2017]. 4 cycles, stoped due to toxicity |
| CO1P0199C KK06 | 47 | F | Colorectal Liver Metastasis | Chemotherapy 7 Cycles [treatment ended 06/09/2017] |
| C01P0112B KK01 | 48 | M | Colorectal Liver Metastasis | Neoadjuvant Chemotherapy. FOLFIRI 12 Cycles from [01/05/2016 to 28/09/2016] |
| C01P0162B KK06 | 48 | F | Focal Nodular Hyperplasia | None |
| C01P0226A KK01 | 48 | M | Gallbladder Adenocarcinoma | None |
| C01P0216B KK06 | 50 | F | Colorectal Liver Metastasis | 8 cycles of CAPOX |
| C01P0261B KK09 | 50 | M | Benign Liver Lesion (originally thought to be potential metastasis from renal carcinoma) | Sorafenib (for renal carcinoma) |
| C01P0032B KK06 | 52 | M | Colorectal Liver Metastasis | Neoadjuvant long course chemoradiatition [Nov 2014], FOLFOX + Cetuximab [01/06/2015-04/09/2015] |
| L8 | 56 | M | Cancer | No data available |
| C01P0126B KK01 | 57 | M | Colorectal Liver Metastasis | Neoadjuvant Chemotherapy. CAPOX 7 Cycles [01/08/2016 to 12/12/2016] |
| C01P0212B KK06 | 58 | F | Colorectal Liver Metastasis | FOLFOX 6 cycles [25/08/2017 to 01/10/2017] |
| C01P0217D KK06 | 58 | F | Colorectal Liver Metastasis | XELOX prior to liver resection [7 cycles ], Long course neoadjuvant chemoradiotherapy |
| C01P0012C KK01 | 62 | M | Colorectal Liver Metastasis | Yes, SCOT trial (OxGdM or XELOX) [01/07/2013 to 30/09/2013] |
| C01P0201B KK06 | 62 | F | Colorectal Liver Metastasis | FOLFOX [30/10/2013-30/03/2014], FOLFIRI (aflibercept added 16/06/2014) [15/04/2014-01/05/2015], FOLFIRI (switched to single agent Irinotecan in March 2016 due to possible angina) [01/09/2015-01/07/2017] |
| C01P0193B KK06 | 64 | F | Vaginal Squamous Cell Carcinoma Liver metastasis | radiotherapy only. |
| C01P0306B KK06 | 64 | M | Colorectal Liver Metastasis | XELOX 3 Cycles [November-December 2018], FOLFOX with oxiplatin added later [December 2018 to February 2019] |
| C01P0300A KK01 | 64 | M | Colorectal Liver Metastasis | FOLFOX + Panitumab, 8 cycles [June-November 2018] |
| C01P0241B KK06 | 65 | M | Colorectal Liver Metastasis | None |
| C01P0124A KK01 | 67 | M | Colorectal Liver Metastasis | Xelox (Oxiplatin stopped after 6 cycles) [Nov 2015 to May 2016], Chemoradiation [June-July 2016]. |
| C01P0131D KK06 | 68 | M | Colorectal Liver Metastasis (also has myelodysplastic syndrome (MDS)) | Azacytidine for MDS [14/09/2015-Jan 2018], only received radiotherapy for colorectal liver mets |
| C01P0058B KK01 | 69 | M | Colorectal Liver Metastasis | None |
| L9 | 69 | M | No data available | No data available |
| C01P0229B KK06 | 70 | M | Colorectal Liver Metastasis | CAPOX [no further details] |
| C01P0075B KK04 | 71 | M | Liver Abscess | Yes |
| L4 | 71 | M | No data available | No data available |
| C01P0103A KK05 | 72 | F | Colorectal Liver Metastasis | Yes, [April-September 2016] |
| C01P0103A KK05 | 72 | F | Colorectal Liver Metastasis | Yes, [April-September 2016] |
| C01P0103A KK05 | 72 | F | Colorectal Liver Metastasis | Yes, [April-September 2016] |
| C01P0234E KK06 | 72 | F | Uveal Melanoma Liver Metastasis | None (Radiotherapy) |
| C01P0249C KK06 | 72 | F | Colorectal Liver Metastasis | Oxaliplatin & capecitabine. Cap converted to Raltitrexed in Jan 2018 and Ox reduced to 75% due to chest pains [Dec 2017 - Feb 2018] |
| C01P0251C KK06 | 72 | M | Colorectal Liver Metastasis | Chemotherapy [treatment ended October 2018] |
| C01P0301A KK06 | 72 | M | Colorectal Liver Metastasis | XELOX 4 cycles [May-July 2018], Cetuximab-FOLFIRI 8 cycles [August-December] |
| C01P0247C KK06 | 73 | M | Colorectal Liver Metastasis | FOLFOX 6 cycles [01/06/2017 - 01/09/2017], Long course neoadjuvant chemoradiation [15/10/2016 - 24/11/2016] |
| C01P0001B KK01 | 74 | M | Colorectal Liver Metastases | None |
| C01P0242B KK06 | 74 | M | Uveal Melanoma Liver Metastasis | None |
| C01P0055B KK05 | 75 | M | Colorectal Liver Metastasis | Oxalaplatin and Capecitabine [29/02/2016-03/05/2016] |
| C01P0330B KK06 | 75 | M | Colorectal Liver Metastasis | Capecitabine [6 months 2016] |
| C01P0108A KK05 | 82 | M | Colorectal Liver Metastasis | CAPOX [No further details] |
| C01P0207A KK06 | 88 | M | Colorectal Liver Metastasis | Capecitabine [01/10/2016-20/03/2017] |
| C01P0096B KK02 | 89 | F | Colorectal Liver Metastasis | None |
| TB15 3915 | 37 | M | Colorectal Liver Metastasis | Xelox and DXT |
| TB16 2481 | 49 | M | Colorectal Liver Metastasis | nil |
| TB16 2721 | 36 | M | Colorectal Liver Metastasis | nil |
| 1PD | | | No data available | No data available |
| 2PD | | | No data available | No data available |
| 4PD | | | No data available | No data available |

### Supplementary Table 2 - Primers

| First Round Primers | | | |
|---|---|---|---|
| **Name** | **5' --->3'** | **Amplicon Region** | **Length (bp)** |
| **AF** | GCTCACATCACCCCATAAAC | 627-646 | 2460 |
| **AR** | GATTACTCCGGTCTGAACTC | 3087-3068 | |
| **BF** | ACCAACAAGTCATTATTACCC | 2395-2415 | 2258 |
| **BR** | TGAGGAAATACTTGATGGCAG | 4653-4633 | |
| **CF** | CCGTCATCTACTCTACCATC | 4489-4508 | 1979 |
| **CR** | GGACGGATCAGACGAAGAG | 6468-6450 | |
| **DF** | AATACCCATCATAATCGGAGG | 6113-6133 | 2324 |
| **DR** | GGTGATGAGGAATAGTGTAAG | 8437-8417 | |
| **EF** | AACCACTTTCACCGCTACAC | 8128-8147 | 2388 |
| **ER** | AGTGAGATGGTAAATGCTAG | 10516-10487 | |
| **FF** | ACTTCACGTCATTATTGGCTC | 9821-9841 | 2280 |
| **FR** | ATAGGAGGAGAATGGGGGATAG | 12101-12080 | |
| **GF** | ACCCCCCACTATTAACCTACTG | 11866-11887 | 2058 |
| **GR** | GGTAGAATCCGAGTATGTTGG | 13924-13904 | |
| **HF** | TATTCGCAGGATTTCTCATTAC | 13721-13742 | 2276 |
| **HR** | AGCTTTGGGTGCTAATGGTG | 15997-15978 | |
| **IF** | CCCATCCTCCATATATCCAAAC | 15659-15680 | 1778 |
| **IR** | GGTTAGTATAGCTTAGTTAAAC | 868-847 | |

| Second Rounds Primers | | | |
|---|---|---|---|
| **A** | **5' --->3'** | **Amplicon Region** | **Length (bp)** |
| **1F** | TGTAAAACGACGGCCAGTTCACCCTCTAAATCACCAG | 721-740 | 547 |
| **1R** | CAGGAAACAGCTATGACCGATGGCGGTATATAGGCTGAG | 1268-1248 | |
| **2F** | TGTAAAACGACGGCCAGTTTAAAACTCAAAGGACCTGGC | 1157-1177 | 552 |
| **2R** | CAGGAAACAGCTcATGACCCTGGTAGTAAGGTGGAGTGGG | 1709-1689 | |
| **3F** | TGTAAAACGACGGCCAGTAACTTAACTTGACCGCTCTGAG | 1650-1671 | 543 |
| **3R** | CAGGAAACAGCTATGACCATTGGTGGCTGCTTTTAGG | 2193-2175 | |
| **4F** | TGTAAAACGACGGCCAGTACTGTTAGTCCAAAGAGGAAC | 2091-2111 | 553 |
| **4R** | CAGGAAACAGCTATGACCTCGTGGAGCCATTCATACAG | 2644-2625 | |
| **B** | **5' --->3'** | **Amplicon Region** | **Length (bp)** |
| **5F** | TGTAAAACGACGGCCAGTCAGTGACACATGTTTAACGGC | 2549-2569 | 538 |
| **5R** | CAGGAAACAGCTATGACCGATTACTCCGGTCTGAACTC | 3087-3068 | |
| **6F** | TGTAAAACGACGGCCAGTCAGCCGCTATTAAAGGTTCG | 3017-3036 | 357 |
| **6R** | CAGGAAACAGCTATGACCGGAGGGGGGTTCATAGTAG | 3374-3356 | |
| **7F** | TGTAAAACGACGGCCAGTCCTTAGCTCTCACCATCGC | 3533-3351 | 524 |
| **7R** | CAGGAAACAGCTATGACCAGAGTGCGTCATATGTTGTTC | 4057-4037 | |
| **8F** | TGTAAAACGACGGCCAGTAATAAACACCCTCACCACTAC | 4005-4025 | 572 |
| **8R** | CAGGAAACAGCTATGACCGTTTATTTCTAGGCCTACTCAG | 4577-4556 | |
| **C** | **5' --->3'** | **Amplicon Region** | **Length (bp)** |
| **9F** | TGTAAAACGACGGCCAGTACACTCATCACAGCGCTAAG | 4518-4537 | 485 |
| **9R** | CAGGAAACAGCTATGACCGATTTTGCGTAGCTGGGTTTG | 5003-4983 | |
| **10F** | TGTAAAACGACGGCCAGTTCCATCATAGCAGGCAGTTG | 4950-4969 | 531 |
| **10R** | CAGGAAACAGCTATGACCTGTAGGAGTAGCGTGGTAAGG | 5481-5462 | |
| **11F** | TGTAAAACGACGGCCAGTACCTCAATCACACTACTCCC | 5367-5386 | 557 |
| **11R** | CAGGAAACAGCTATGACCTAGTCAACGGTCGGCGAAC | 5924-5906 | |
| **12F** | TGTAAAACGACGGCCAGTCACTCAGCCATTTTACCTCAC | 5875-5895 | [illegible] |

| | | | |
|---|---|---|---|
| **12R** | CAGGAAACAGCTATGACCATGGCAGGGGGTTTTATATTG | 6430-6410 | [illegible] |
| **D** | **5' --->3'** | **Amplicon Region** | **Length (bp)** |
| **13F** | TGTAAAACGACGGCCAGTTTAGGGGCCATCAATTTCATC | 6378-6398 | 566 |
| **13R** | CAGGAAACAGCTATGACCAAGAAAGATGAATCCTAGGGC | 6944-6924 | |
| **14F** | TGTAAAACGACGGCCAGTATTTAGCTGACTCGCCACAC | 6863-6882 | 533 |
| **14R** | CAGGAAACAGCTATGACCCATCCATATAGTCACTCCAGG | 7396-7376 | |
| **15F** | TGTAAAACGACGGCCAGTGGCTCATTCATTTCTCTAACAG | 7272-7293 | 519 |
| **15R** | CAGGAAACAGCTATGACCGGCAGGATAGTTCAGACGG | 7791-7773 | |
| **16F** | TGTAAAACGACGGCCAGTTAACATCTCAGACGCTCAGG | 7744-7763 | 557 |
| **16R** | CAGGAAACAGCTATGACCTACAGTGGGCTCTAGAGGG | 8301-8283 | |
| **E** | **5' --->3'** | **Amplicon Region** | **Length (bp)** |
| **17F** | TGTAAAACGACGGCCAGTACAGTTTCATGCCCATCGTC | 8196-8215 | 544 |
| **17R** | CAGGAAACAGCTATGACCGTATAAGAGATCAGGTTCGTC | 8740-8720 | |
| **18F** | TGTAAAACGACGGCCAGTACCACCCAACAATGACTAATC | 8656-8676 | 545 |
| **18R** | CAGGAAACAGCTATGACCGTTGTCGTGCAGGTAGAGG | 9201-9183 | |
| **19F** | TGTAAAACGACGGCCAGTATCCTAGAAATCGCTGTCGC | 9127-9146 | 540 |
| **19R** | CAGGAAACAGCTATGACCATTAGACTATGGTGAGCTCAG | 9661-9641 | |
| **20F** | TGTAAAACGACGGCCAGTCATCCGTATTACTCGCATCAG | 9607-9627 | 540 |
| **20R** | CAGGAAACAGCTATGACCTAGCCGTTGAGTTGTGGTAG | 10147-10128 | |
| **F** | **5' --->3'** | **Amplicon Region** | **Length (bp)** |
| **21F** | TGTAAAACGACGGCCAGTCAACACCCTCCTAGCCTTAC | 10085-10104 | 564 |
| **21R** | CAGGAAACAGCTATGACCAGGCACAATATTGGCTAAGAG | 10649-10629 | |
| **22F** | TGTAAAACGACGGCCAGTATCGCTCACACCTCATATCC | 10534-10553 | 575 |
| **22R** | CAGGAAACAGCTATGACCATGATTAGTTCTGTGGCTGTG | 11109-11089 | |
| **23F** | TGTAAAACGACGGCCAGTCTAATCTCCCTACAAATCTCC | 11054-11074 | 551 |
| **23R** | CAGGAAACAGCTATGACCTAGGTCTGTTTGTCGTAGGC | 11605-11586 | |
| **24F** | TGTAAAACGACGGCCAGTTCCTTGTACTATCCCTATGAG | 11541-11561 | 513 |
| **24R** | CAGGAAACAGCTATGACCCGTGTGAATGAGGGTTTTATG | 12054-12034 | |
| **G** | **5' --->3'** | **Amplicon Region** | **Length (bp)** |
| **25F** | TGTAAAACGACGGCCAGTACAATGGGGCTCACTCACC | 12001-12019 | 544 |
| **25R** | CAGGAAACAGCTATGACCGTGGCTCAGTGTCAGTTCG | 12545-12527 | |
| **26F** | TGTAAAACGACGGCCAGTCATGTGCCTAGACCAAGAAG | 12498-12517 | 511 |
| **26R** | CAGGAAACAGCTATGACCCTGATTTGCCTGCTGCTGC | 13009-12991 | |
| **27F** | TGTAAAACGACGGCCAGTGCCCTTCTAAACGCTAATCC | 12940-12959 | 513 |
| **27R** | CAGGAAACAGCTATGACCGGGAGGTTGAAGTGAGAGG | 13453-13435 | |
| **28F** | TGTAAAACGACGGCCAGTCGGGTCCATCATCCACAAC | 13365-13383 | 494 |
| **28R** | CAGGAAACAGCTATGACCGTTAGGTAGTTGAGGTCTAGG | 13859-13839 | |
| **H** | **5' --->3'** | **Amplicon Region** | **Length (bp)** |
| **29F** | TGTAAAACGACGGCCAGTACCTAAAACTCACAGCCCTC | 13790-13809 | 584 |
| **29R** | CAGGAAACAGCTATGACCAGGATTGGTGCTGTGGGTG | 14374-14356 | |
| **30F** | TGTAAAACGACGGCCAGTCAACCACCACCCCATCATAC | 14331-14350 | 526 |
| **30R** | CAGGAAACAGCTATGACCAAGGAGTGAGCCGAAGTTTC | 14857-14838 | |
| **31F** | TGTAAAACGACGGCCAGTATTCATCGACCTCCCCACC | 14797-14815 | 571 |
| **31R** | CAGGAAACAGCTATGACCGGTTGTTTGATCCCGTTTCG | 15368-15349 | |
| **32F** | TGTAAAACGACGGCCAGTAGCCCTAGCAACACTCCAC | 15316-15334 | 580 |
| **32R** | CAGGAAACAGCTATGACCTACAAGGACAGGCCCATTTG | 15896-15877 | |
| **I** | **5' --->3'** | **Amplicon Region** | **Length (bp)** |
| **D1F** | TGTAAAACGACGGCCAGTATCGGAGGACAACCAGTAAG | 15758-15777 | 536 |
| **D1R** | CAGGAAACAGCTATGACCGTGGGTAGGTTTGTTGGTATC | 16294-16274 | |
| **D2F** | TGTAAAACGACGGCCAGTCTCAACTATCACACATCAACTG | 16223-16244 | [illegible] |

| | | | |
|---|---|---|---|
| **D2R** | CAGGAAACAGCTATGACCAGATACTGCGACATAGGGTG | 129-110 | [illegible] |
| **D3F** | TGTAAAACGACGGCCAGTCACCCTATTAACCACTCACG | 15-34 | 374 |
| **D3R** | CAGGAAACAGCTATGACCCTGGTTAGGCTGGTGTTAGG | 389-370 | |
| **D4F** | TGTAAAACGACGGCCAGTGCCACAGCACTTAAACACATC | 323-343 | 448 |
| **D4R** | CAGGAAACAGCTATGACCTGCTGCGTGCTTGATGCTTG | 771-752 | |

**Supplementary Methods**

**Methods S1**

### 2-dimensional spatial simulations and Bayesian inference

#### S1.1 Model outline

Using the Gillespie algorithm, we simulate stochastic cell division, death and mtDNA mutation on a 2-dimensional lattice to explore the effects of a periportal progenitor niche on the patterns of mtDNA mutations. In simulations, cells are characterised by 3 quantities: their x- and y-coordinates on the lattice, and a list of mutations that they carry.

Cells acquire new mtDNA mutations via two processes. First, mutations which accumulate in mtDNA as a result of normal mitochondria turnover throughout the cell cycle are modelled by means of a third process in the Gillespie algorithm: occurring independently alongside cell division and death. We assume that, on average, each mitochondrion is turned over once per cell division. Secondly, cell divisions will lead to increased mutation accumulation in the mitochondria due to cells replenishing their mitochondria copy number following mitosis. Accordingly, alongside inheriting all mutations carried by the parent cell, daughter cells may also acquire new mtDNA mutations immediately after cell division. We estimate the mutation rate in mitochondria, $\mu$, to be equal to 0.1 per division across the whole mitochondrial genome, based on recent estimations of the mutation rate in healthy nuclear DNA (1) and estimations of the mutation rate in mitochondrial DNA versus nuclear DNA (2).

To model the displacement of neighbouring cells during division, we adapted an algorithm developed by Waclaw et al. (3). In our simulations we extend these mechanics by accounting for relative positions of cells during pushing, favouring straight-line pushing over displacement in other directions, in accordance with Newton’s second law of motion.

The simulated system is mostly comprised of hepatocyte-like ‘non-PT’ cells, and a small number, $N_{PT}$, of progenitor-like ‘PT cells’. PT and non-PT cells differ by their replication rates, which are linearly related through the scalar model parameter $\beta$ (see main Methods), and by their location on the lattice. Non-PT cells occupy unique, discretised coordinates on the lattice, whereas PT cells are modelled as a well-mixed pool of size $N_{PT}$. PT cells couple to the lattice through a single cell situated at the centre of the system, which acts as a proxy for all cells in the PT cell pool. When a PT cell divides, one daughter cell remains in the PT cell pool, whilst the other becomes a non-PT cell and enters the lattice adjacent to the proxy cell, pushing neighbouring cells outwards if necessary. PT cells divide strictly asymmetrically (i.e. by producing one PT and one non-PT cell) in order to maintain a constant number of PT cells in the system.

#### S1.2 Spatial simulations of mtDNA dynamics

A single simulation is split into two main phases. During the first phase, a single cell is seeded in the centre of the lattice, which then divides (without any mutations occurring) until the system is comprised of $N_{max}$ cells with zero mutations. With the system initialised, the simulation enters into the second phase. mtDNA mutation is switched on, and successive "rounds" of homeostasis are simulated. In each round, first a pre-specified fraction of non-PT cells is killed at random and then the remaining cells divide (without any cell death) until the system size returns to $N_{max}$ cells. Once the total number of cells has reached $N_{max}$, the round has finished and the system is measured. When taking measurements of the spatial data, only cells located in the lattice are recorded, i.e. mutations in the PT cells are not counted. This method of measurement is reasonable since, when analysing the tissue data, the sections in general do not represent the entirety of the clonal patch of CCO- cells along the portal tract to central vein axis. A single simulation is summarised in the pseudo-code present in Algorithm 1.

Full code is available at

https://github.com/MagnusHaughey/HepatocyteClonalDynamics

#### S1.3 Bayesian inference to compare model hypothesis with multi-region genomic data

To enable comparison of simulation data to the multi-region sampled data, we employ a zonal random sampling method to the spatial data, in which we divide the cells into different sections based on their radial distance from the centre of the system. We sample $N_S$ cells from each section and measure the variety of mtDNA mutations carried by cells in each section. We adopt the convention of naming the central section (in which the PT cells reside) as “section 1” with sections 2, 3 etc. located at increasing radial distances from the centre of the system. In the same fashion as the NGS analysis, we analysed the total number of different mtDNA mutations detected within each sampled section. The same mutation carried by two or more cells is not counted more than once. Since our measure of mutational burden reflects the level of genetic diversity in the sampled cells, this value will have some dependence on sample size, $N_S$, per section. In the experimental data, we estimate this based on our cut size of 6,500 $\mu m^2$ across 10 tissue sections, each at a section thickness of 10μm. The volume of each cut, $V_{cut}$, is then approximately equal to:

$$V_{cut} \approx 6,500\ \mu m^2 \times 10\ \mu m \times 10 = 650,000 \mu m^3.$$

Estimated values for hepatocyte volume, $V_{hep}$, vary in the literature, ranging from ≈ 3,400 $\mu m^3$ to 12,000 $\mu m^3$ (4-6), with some studies observing non-constant distributions of hepatocyte volume along the portal tract to central vein axis. Assuming a constant value for hepatocyte volume and taking the upper bound of this range, $V_{hep}$ = 12,000 $\mu m^3$, leads to an estimated sample size of approximately 60 cells per cut. This value is consistent with the number of cells counted visually from the H&E slides. As such, we then also sample $N_S$ = 60 cells from each section in our simulated data.

We implemented an Approximate Bayesian Computation (ABC) rejection sampling algorithm to estimate the joint posterior distribution of model parameters β and $N_{PT}$, obeying the following steps:

1. Sample parameters $\beta^* \sim U(0.1, 50.0)$ and $N^*_{PT} \sim U(10, 200)$.
2. Simulate with $\beta^*$ and $N^*_{PT}$ for a total of T=100 rounds of homeostasis.
3. Accept $\beta^*$ and $N^*_{PT}$ based on the squared Euclidean distance between experimental and simulated data sets.
4. Repeat.

As indicated in step 2, we do not infer the number of rounds of homeostasis, T. Instead, we perform a grid-search i.e. for each sampled pair of $\beta^*$ and $N^*_{PT}$ we evolve the system for a total of T = 100 rounds. After each round we compare the simulated data (by zonal sampling in 5 random radial directions) to the experimental data. If any of the 5 individual measurements are sufficiently close to the sample data then we accept $\beta^*$ and $N^*_{PT}$ and the round number, *t*. After simulating with a sufficient number of $\beta^*$ and $N^*_{PT}$ parameter pairs we then select the round number, *t*, with the greatest number of accepted parameter pairs and plot the distribution of accepted $\beta^*$ and $N^*_{PT}$ corresponding to this value of *t*. We then summarised these distributions using geometric median as a point estimator for $\beta^*$ and $N^*_{PT}$. For each experimental sample, we set the required squared Euclidean distance between simulated and experimental data such that the algorithm has a 0.1% acceptance rate.

**Supplementary Methods S2**

### 1-dimensional liver expansion dynamics simulations

#### S2.1 Model outline

We simulate 1D clonal expansion of hepatocytes, exploring the effect of slow or rapid clonal expansion on the number of shared mtDNA mutations between spatially adjacent sampled regions. In addition to slow and rapid expansion, we compare the impact of the expanded patch of cells occupying a quiescent and "streaming" state for specified periods of time. A summary of the simulations follows.

Simulations begin with a single PT cell seeded on the lattice. This PT cell has the same properties as those in the 2D simulations, *i.e.* it divides strictly asymmetrically (divides into 1 PT cell and 1 hepatocyte), therefore maintaining a constant number of PT cells in the system, and it can never die. Furthermore, in these 1D simulations the PT cell strictly only divides in one direction (*i.e.* can not produce a hepatocyte on either its left or right, but only in one direction).

Since we are only interested in somatic mutations that are acquired during the clonal expansion of the PT cell, the PT cell begins the simulation carrying zero mutations. Subsequently, we simulate cell division and mtDNA mutation using the Gillespie algorithm. Hepatocytes die stochastically at a low rate, and can proliferate to replace lost cells if a neighbouring hepatocyte dies. Empty spaces left behind by dying hepatocytes can also be filled if neighbouring hepatocytes are pushed into the space due to the PT cell proliferating and producing a new hepatocyte.

mtDNA dynamics are modelled in the same fashion as in the 2D simulations. It is assumed that mtDNA carried by the PT cell and all hepatocytes mutate at the same rate of $5\times10^{-5}$ mutations per mtDNA per day (7). As in the 2D simulations, cells accumulate mtDNA mutations continuously throughout the cell cycle, and may also mutate further during cell division as a result of the cell replenishing normal mitochondria copy number.

#### S2.2 Simulation of expansion timescales in 1D systems

Simulations consist of two phases. The first phase is the clonal expansion phase, which begins with the PT cell being seeded, and ends when the system reaches a size of $N_{max}$ hepatocytes. Following the clonal expansion phase is a period of either quiescence or "streaming" dynamics which lasts for time $t^*$, where $t^*$ is an input parameter of the model. For quiescent dynamics, all cells (hepatocytes and single PT cell) stop dividing and dying, with the only process to continue being continuous mtDNA mutations. For "streaming" dynamics the hepatocytes again stop dividing and dying, however the PT cell continues to divide, and pushes more hepatocytes into the system. At the other end of the system, hepatocytes that are pushed far enough from the PT, such that they exceed a distance of $N_{max}$ lattice points from the PT, are removed from the system. The result of these rules is "conveyor belt" or "streaming" dynamics, with new hepatocytes being generated at the PT end of the system, and hepatocytes being lost at the opposite end.

To measure the resulting system, we adapt the zonal sampling method used to measure the 2D system. In this 1D system, after the second phase (period of either quiescent or "streaming" dynamics) has ended we separate the system into five sections, each containing an equal number of cells. Within each section, we pool the mutational data for all cells and measure the variety of SNVs within the section. We then compare the spectrum of non-public mtDNA mutations (mutations not detected in all sections) within each pair of neighbouring sections (*i.e.* compare section 1 to section 2; section 2 to section 3 etc.) and compute the number of mutations which are not shared between both sections (we refer to these as "unique" mutations) as a fraction of the total number of mutations detected in both sections,

$$\frac{|U|}{|A \cup B|},$$

where U denotes the set of unique mutations across both sections, A and B denote the set of all non-public mutations detected in both sections, and |.| denotes magnitude.

We explore the impact of slow and rapid PT expansion rates, as well as quiescent and "streaming" dynamics, on the fraction of unique mutations across the 1D system (Sup. Fig. 17). In both slow and rapid PT expansion rates, motivated by the results of the ABC optimisation performed using the 2D simulations, the PT divides at a faster rate than the hepatocytes. In the slow PT expansion regime, the PT cell divides at a rate of $100^{-1}$ $days^{-1}$ whereas in the rapid case the PT division rate is $7^{-1}$ $days^{-1}$. In both cases, hepatocytes die at a rate of $300^{-1}$ $days^{-1}$. We simulate the second phase (quiescent or "streaming") for $t^*$ equal to 0 days, 30 days, 5 years and 20 years.

Here we illustrate our simulation as follow and the full code is available at

https://github.com/MagnusHaughey/HepatocyteClonalDynamics

**Stage one:** Initiation

Mutation rate = 0

Seed first cell in lattice centre

**while** $N_{cells} < N_{max}$ **do**

Pick random *x* and *y* co-ordinates

Divide( Cell(x,y) )

**end while**

Seed PT cells in separate vector of length $N_{PT}$

**Stage two:** Simulate dynamics

Mutation rate = $\mu$

**For** *T* rounds, **do**

Kill fraction $f_{kill}$ of cells at random

**while** $N_{cells} < N_{max}$ **do**

Pick random *x* and *y* co-ordinates

Divide( Cell(x,y) )

**end while**

Take measurement of system

**end for**

**Algorithm 1:** 2D spatial simulations of mtDNA dynamics

**Supplementary Table 3:** Model parameters used in 1D simulations of $CCO^-$ patch expansion.

| ***Quantity*** | ***Value*** | ***Notes*** |
|---|---|---|
| *Max. system length, $N_{max}$* | 50 cells | |
| *Hepatocyte turnover rate* | $300^{-1}$ per day | (8), (9) |
| *Slow PT division rate* | $100^{-1}$ per day | Faster rate of division for PT cell versus hepatocyte suggested by 2D modelling |
| *Rapid PT division rate* | $7^{-1}$ per day | |
| *mtDNA copy number* | 1,000 per cell | Based on measurements of mtDNA copy number |
| *mtDNA mutation rate* | $5\times10^{-5}$ mutations per mtDNA per day | (7) |
| *Phase 2 length, $t^*$* | { 0 days, 1 month, 5 years, 20 years } | |

### Supplementary Figures:

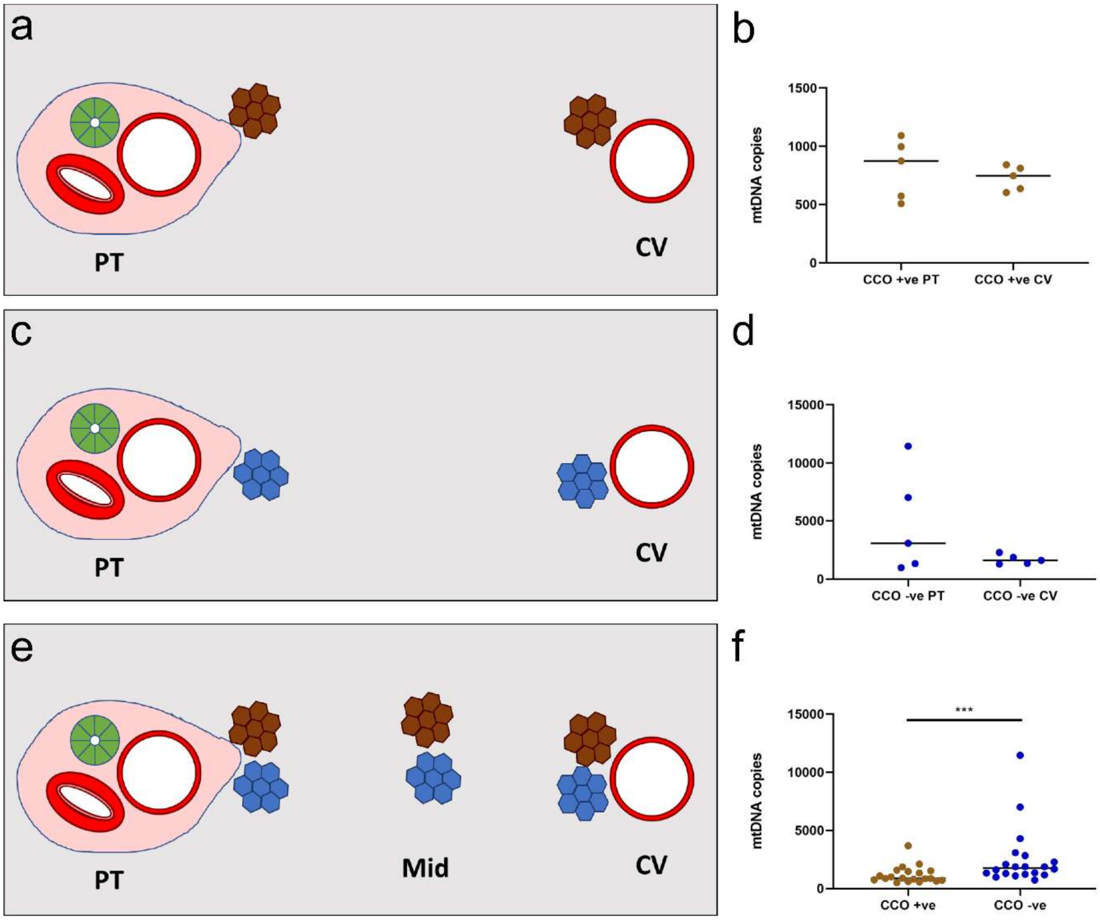


**Figure S1** mtDNA copy number comparisons across the lobule. **(a-d)** mtDNA copy numbers in hepatocytes adjacent to portal tracts (PT) and central veins (CV) from both CCO+ **(a&b)** and CCO- **(c&d)** cells, n=5 samples. **(e&f)** mtDNA copy number comparison between CCO+ and CCO- hepatocytes microdissected from portal, midzonal and pericentral locations, n=20 samples. Bars represent medians and *P****<0.001.

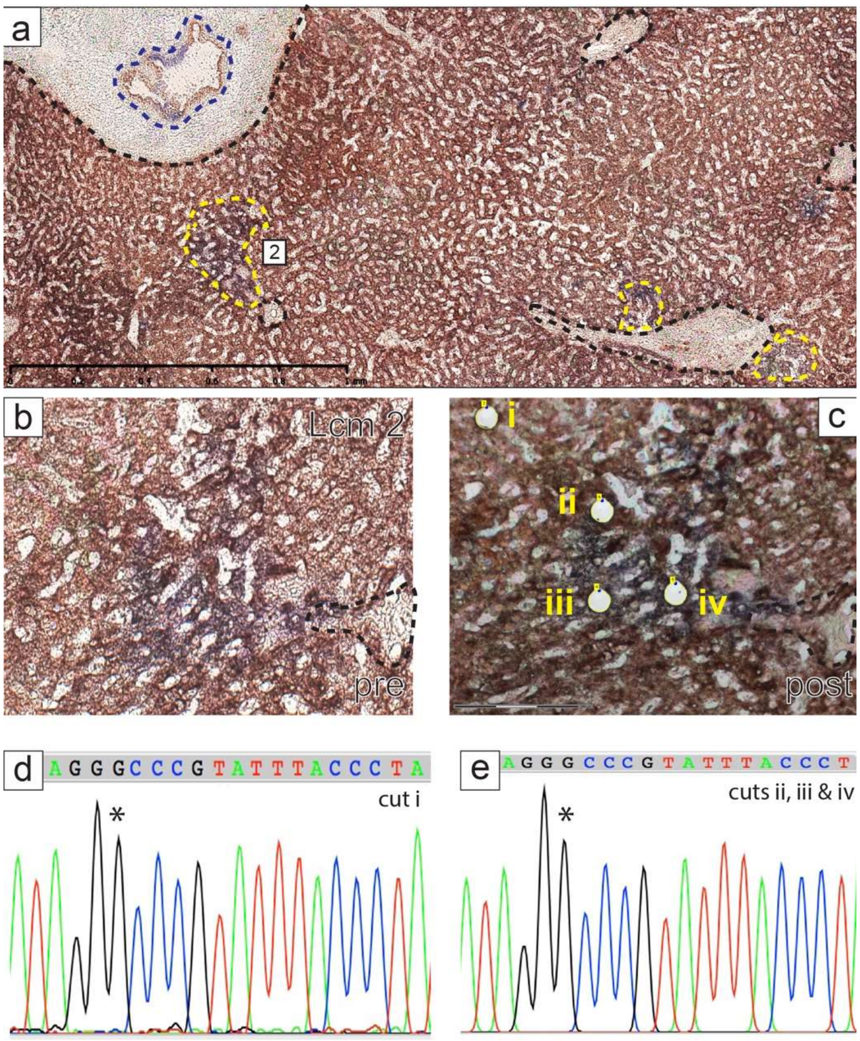


**Figure S2.** Clonal relationships in LCM region 2. **(a)** A CCO/SDH-stained section highlighting a portal tract with biliary epithelium that is partly CCO-deficient and nearby CCO-deficient hepatocyte patches. Portal tracts as well as CCO-deficient hepatocyte patches and biliary epithelium are outlined in black, yellow and blue respectively. **(b&c)** High power images of LCM area 2, pre and post microdissection of CCO-deficient hepatocytes respectively (cells labelled i-iv). **(d)** Sanger sequencing electropherogram of a CCO-proficient hepatocyte (i) and **(e)** Electropherogram for **ii-iv** demonstrating it is wild-type for the *m.8251G>A* mutant present in a CCO-deficient hepatocytes described in Figure 2.

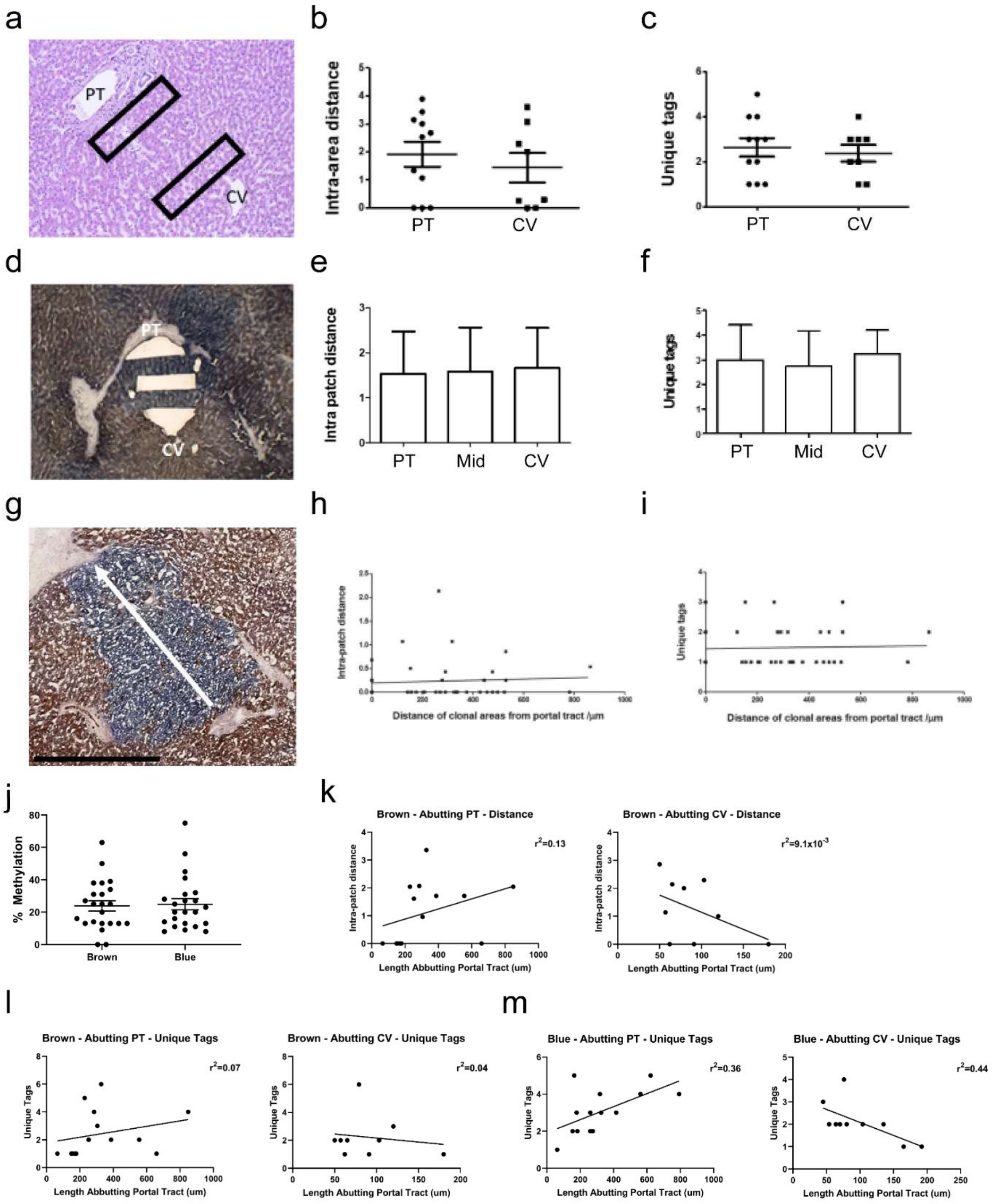
a
PT
CV
b
Intra-area distance
PT
CV
c
Unique tags
PT
CV
d
PT
CV
e
Intra patch distance
PT
Mid
CV
f
Unique tags
PT
Mid
CV
g
h
Intra-patch distance
Distance of clonal areas from portal tract /µm
i
Unique tags
Distance of clonal areas from portal tract /µm
j
% Methylation
Brown
Blue
k
Brown - Abutting PT - Distance
$r^2$=0.13
Intra-patch distance
Length Abbutting Portal Tract (um)
Brown - Abutting CV - Distance
$r^2$=9.1x10$^{-3}$
Intra-patch distance
Length Abutting Portal Tract (um)
l
Brown - Abutting PT - Unique Tags
$r^2$=0.07
Unique Tags
Length Abbutting Portal Tract (um)
Brown - Abutting CV - Unique Tags
$r^2$=0.04
Unique Tags
Length Abutting Portal Tract (um)
m
Blue - Abutting PT - Unique Tags
$r^2$=0.36
Unique Tags
Length Abutting Portal Tract (um)
Blue - Abutting CV - Unique Tags
$r^2$=0.44
Unique Tags
Length Abutting Portal Tract (um)

**Figure S3.** Methylation diversity in normal human liver. **(a)** An H&E with labels denoting large periportal and pericentral areas cut for methylation diversity measurements. **(b&c)** Two measures of methylation diversity: epigenetic distance and unique tags; measured from 11 PT and 8 CV equally-sized (defined per unit area as $mm^2$) tissue extractions as depicted in **(a)**. **(d)** A CCO/SDH stained section displaying PT, midzonal and CV laser microdissections. **(e&f)** Distance and unique tag measurements for equally sized PT, midzonal and CV extractions as depicted in **(d)**. **(g)** A CCO/SDH stained section displaying a large patch stretching from PT to CV. **(h&i)** Epigenetic distance and unique tag diversity measures plotted against the anatomical distance of the tissue extraction from the PT. Extractions were made along the PT-CV axis (arrow in **(g)**) (n=30). **(j)** The percentage of methylated CpG sites within CCO-sufficient (brown) and deficient (blue) microdissections. **(k&l)** Epigenetic distance and unique tag diversity measurements from CCO-sufficient hepatocyte extractions of equal size, abutting 12 PTs and 9 CVs. **(m)** Unique tag measurements from extractions of equal size in 14 PTs and 10 CVs. Graphs in **(k-m)** are plotted against the total length of the CCO-proficient region or CCO-deficient patch that was abutting the PT or CV that each extraction was taken from. Statistical tests performed by Mann-Whitney and Pearson correlation coefficient. No statistical significance observed.

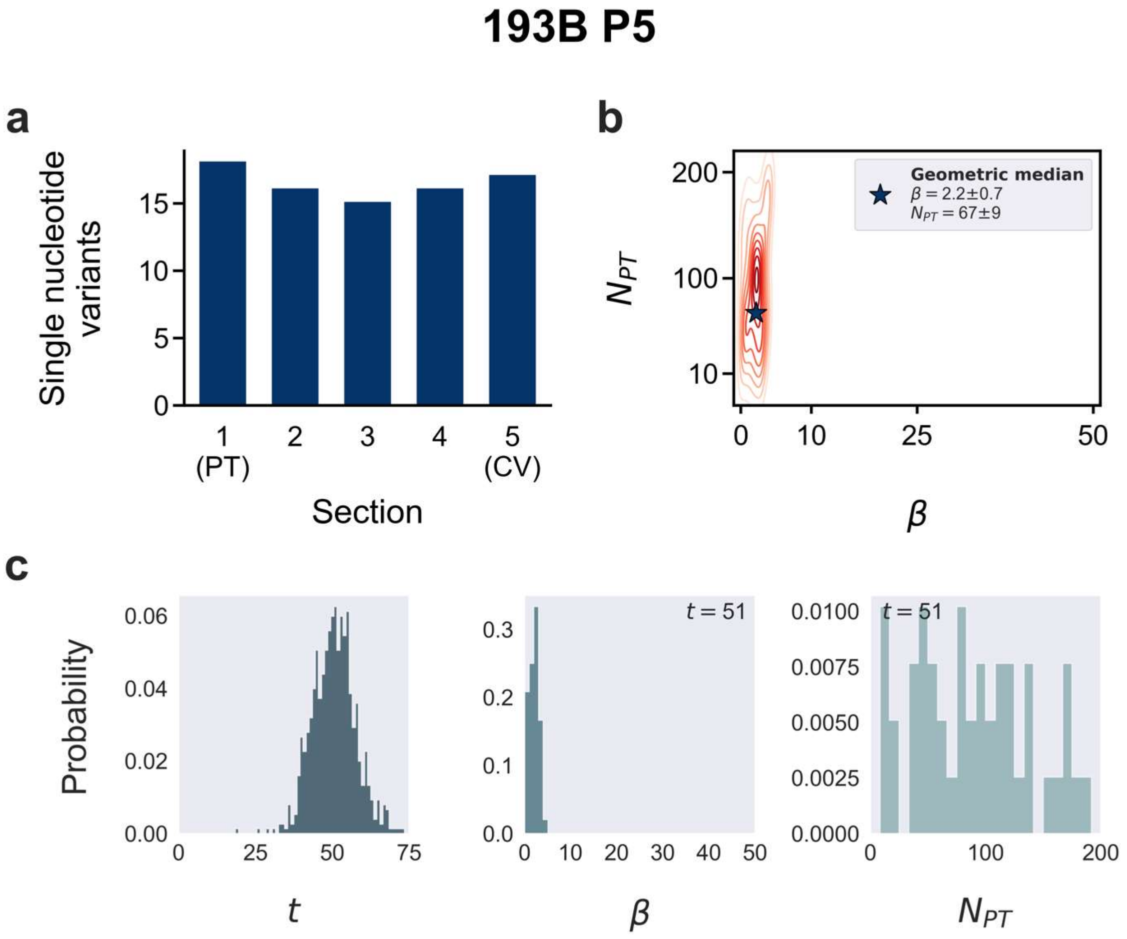


**Figures S4—17.** Individual analyses for each portal-central axis simulated. **(a)** Measured number of unique single nucleotide variants in sequenced tissue. **(b)** Estimated joint posterior distribution of parameters of $\beta$ and $N_{PT}$. Contour lines indicate curves of constant density. A geometric median is used as a point estimator for $\beta$ and $N_{PT}$ and is indicated in the plot as a star. **(c)** Distribution of accepted timepoint, t, $\beta$ and $N_{PT}$ over their respective ranges. $\beta$ and $N_{PT}$ pairs were filtered on the value of the most frequently accepted timepoint (i.e. the highest bin in the left plot, the value of which is indicated in the top corner of middle and right panels).

# L9 Pb

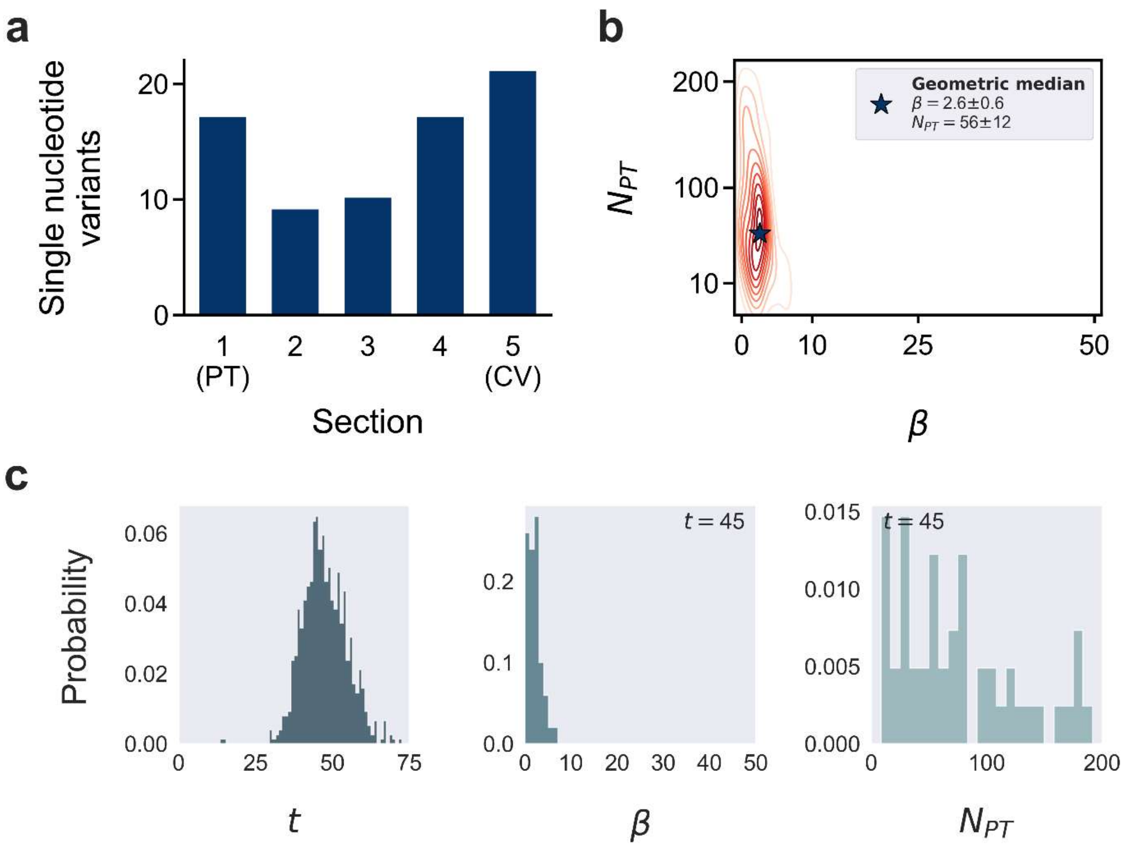
a
Single nucleotide variants
20
10
0
1 (PT)
2
3
4
5 (CV)
Section
b
Geometric median
β = 2.6±0.6
N_PT = 56±12
N_PT
200
100
10
0
10
25
50
β
c
Probability
0.06
0.04
0.02
0.00
0
25
50
75
t
t = 45
0.2
0.1
0.0
0
10
20
30
40
50
β
0.015
t = 45
0.010
0.005
0.000
0
100
200
N_PT

# L9 P12L2

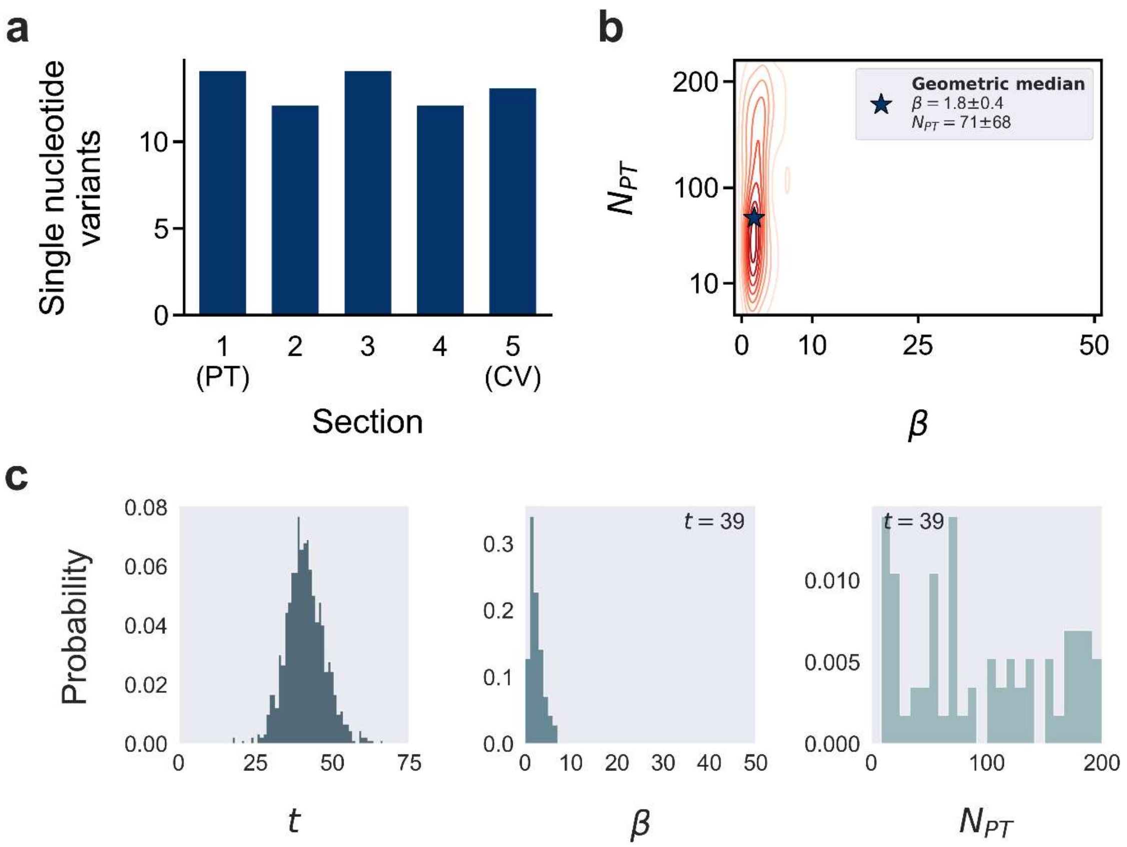
a
Single nucleotide variants
0
5
10
1 (PT)
2
3
4
5 (CV)
Section
b
Geometric median
β = 1.8±0.4
N_PT = 71±68
N_PT
200
100
10
0
10
25
50
β
c
Probability
0.08
0.06
0.04
0.02
0.00
0
25
50
75
t
t = 39
0.3
0.2
0.1
0.0
0
10
20
30
40
50
β
t = 39
0.010
0.005
0.000
0
100
200
N_PT

# L9 P12L1

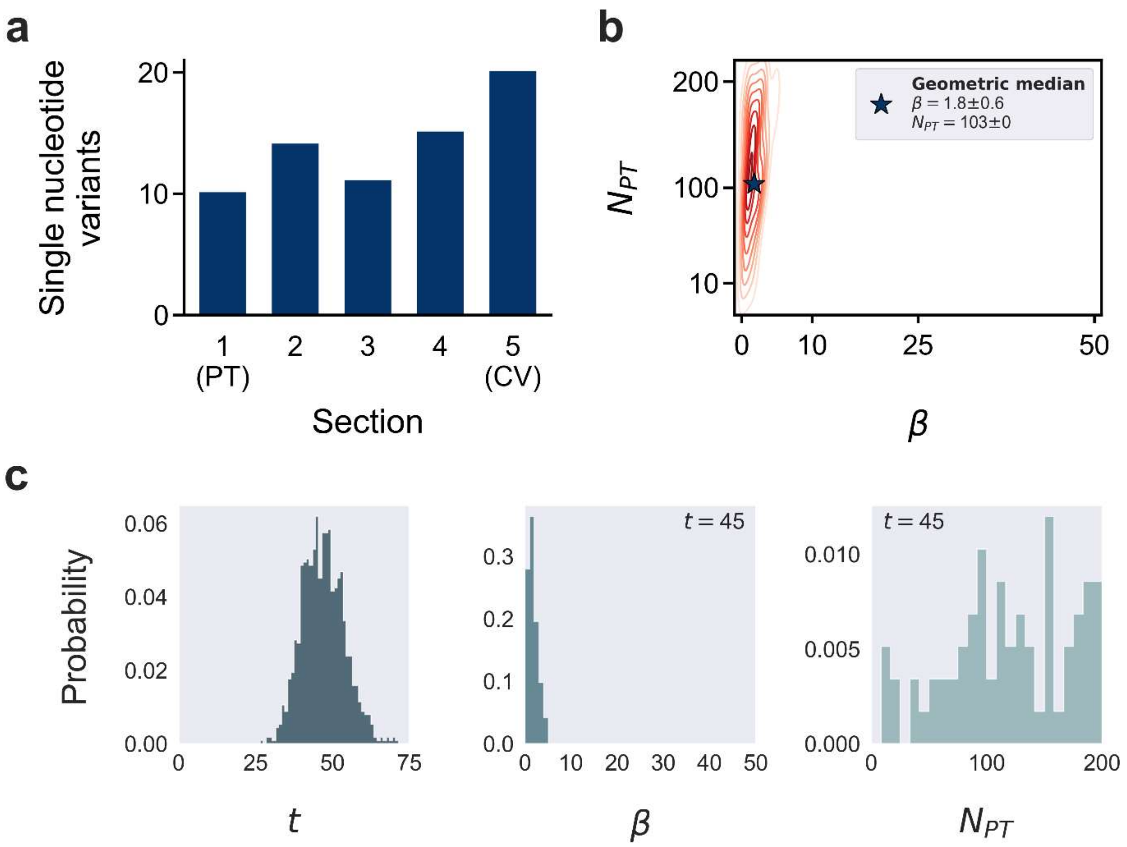
a
Single nucleotide variants
20
10
0
1
(PT)
2
3
4
5
(CV)
Section
b
Geometric median
β = 1.8±0.6
N_PT = 103±0
200
100
10
N_PT
0
10
25
50
β
c
Probability
0.06
0.04
0.02
0.00
0
25
50
75
t
t = 45
0.3
0.2
0.1
0.0
0
10
20
30
40
50
β
t = 45
0.010
0.005
0.000
0
100
200
N_PT

# L9 P1

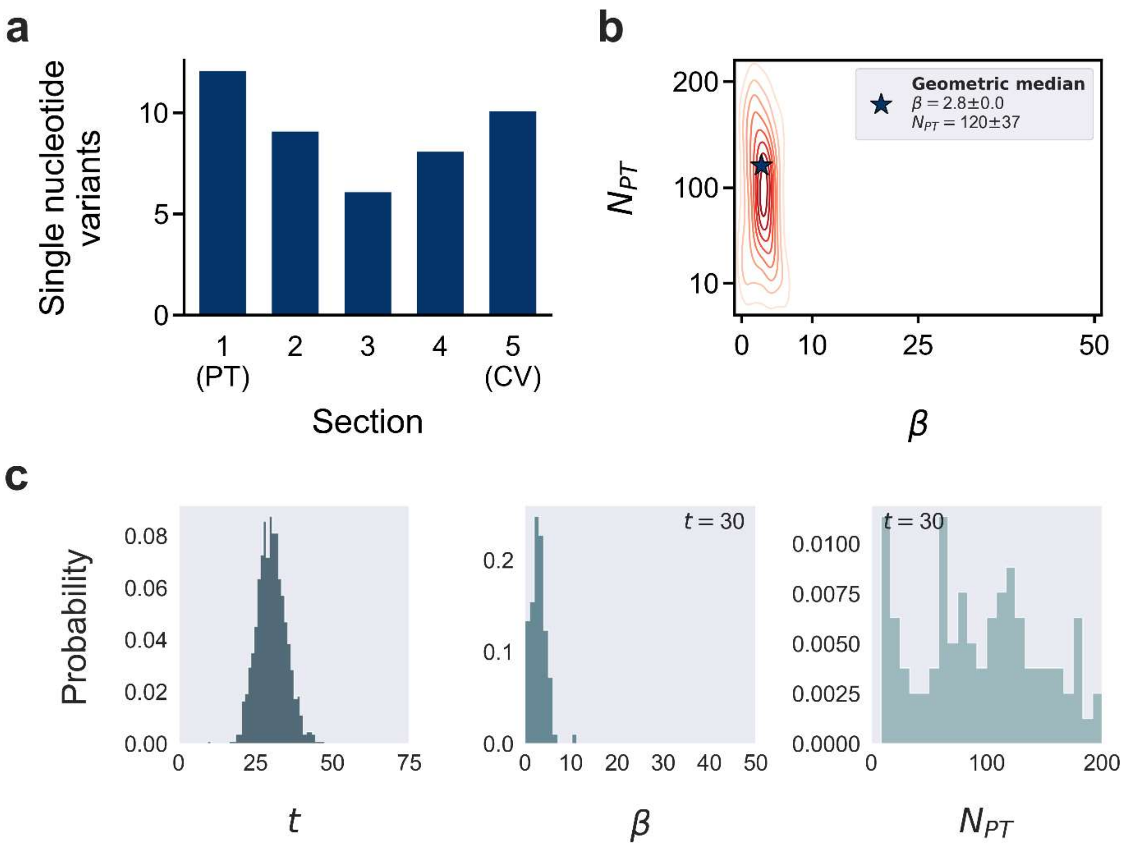
a
Single nucleotide variants
10
5
0
1
(PT)
2
3
4
5
(CV)
Section
b
Geometric median
$\beta = 2.8 \pm 0.0$
$N_{PT} = 120 \pm 37$
$N_{PT}$
200
100
10
0
10
25
50
$\beta$
c
Probability
0.08
0.06
0.04
0.02
0.00
0
25
50
75
$t$
$t = 30$
0.2
0.1
0.0
0
10
20
30
40
50
$\beta$
$t = 30$
0.0100
0.0075
0.0050
0.0025
0.0000
0
100
200
$N_{PT}$

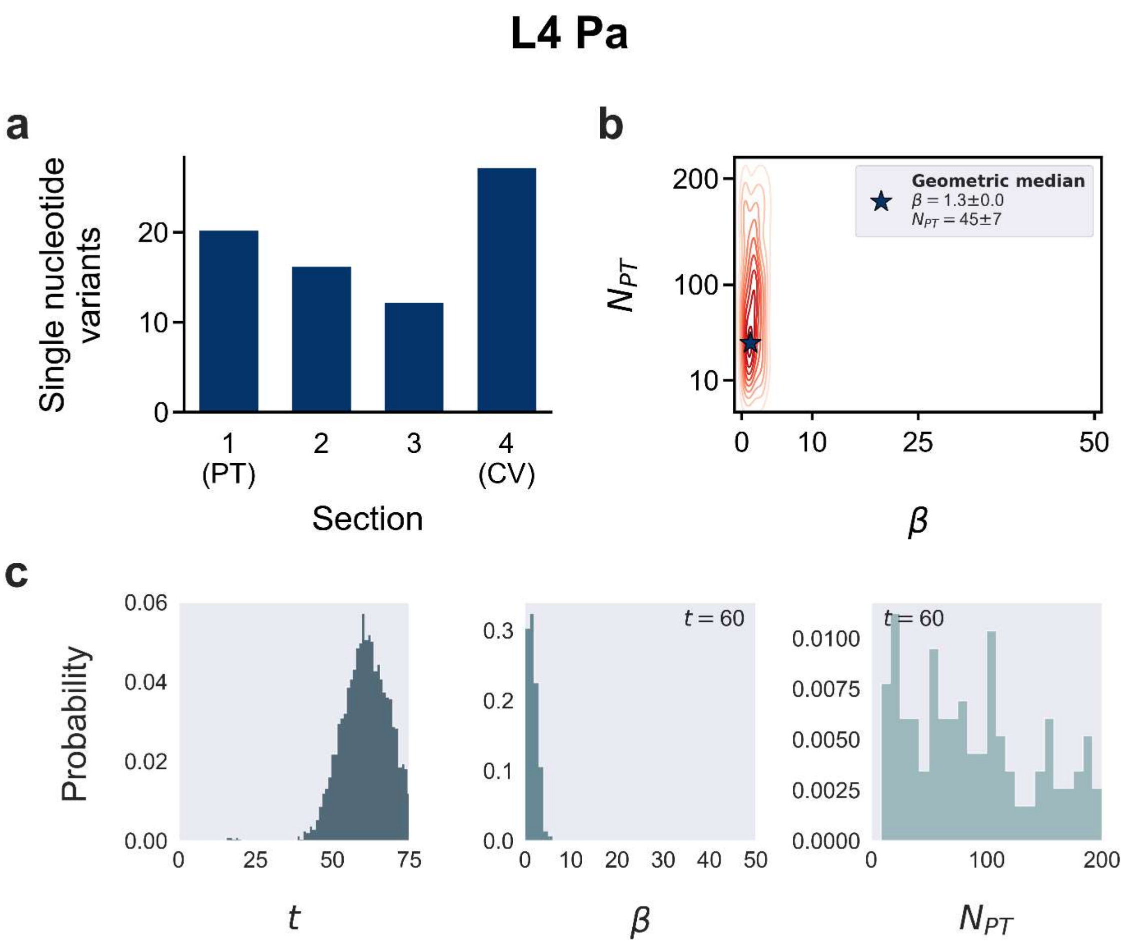
L4 Pa
a
Single nucleotide variants
20
10
0
1
(PT)
2
3
4
(CV)
Section
b
Geometric median
β = 1.3±0.0
N_PT = 45±7
200
100
10
0
10
25
50
β
c
Probability
0.06
0.04
0.02
0.00
0
25
50
75
t
t = 60
0.3
0.2
0.1
0.0
0
10
20
30
40
50
β
t = 60
0.0100
0.0075
0.0050
0.0025
0.0000
0
100
200

# 251B P15

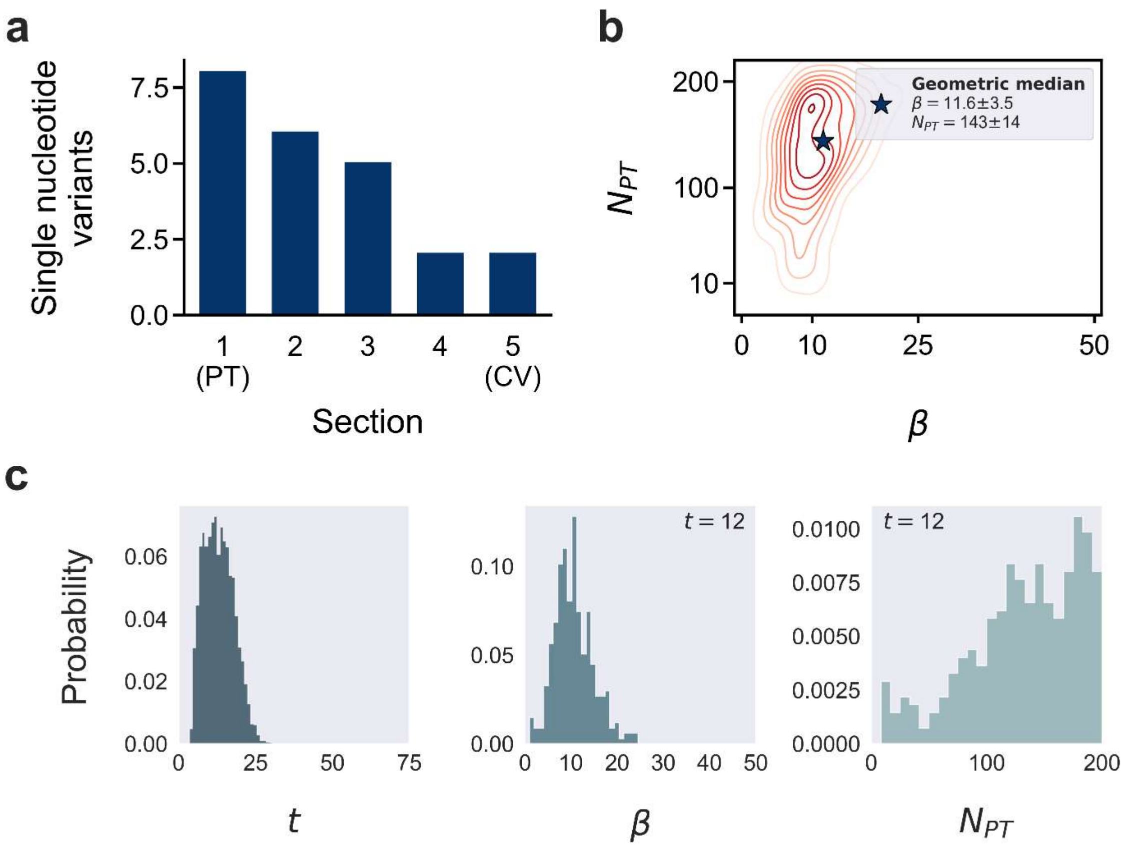
a
Single nucleotide variants
7.5
5.0
2.5
0.0
1
(PT)
2
3
4
5
(CV)
Section
b
Geometric median
β = 11.6±3.5
N_PT = 143±14
200
100
10
N_PT
0
10
25
50
β
c
Probability
0.06
0.04
0.02
0.00
0
25
50
75
t
t = 12
0.10
0.05
0.00
0
10
20
30
40
50
β
t = 12
0.0100
0.0075
0.0050
0.0025
0.0000
0
100
200
N_PT

# 201B P9

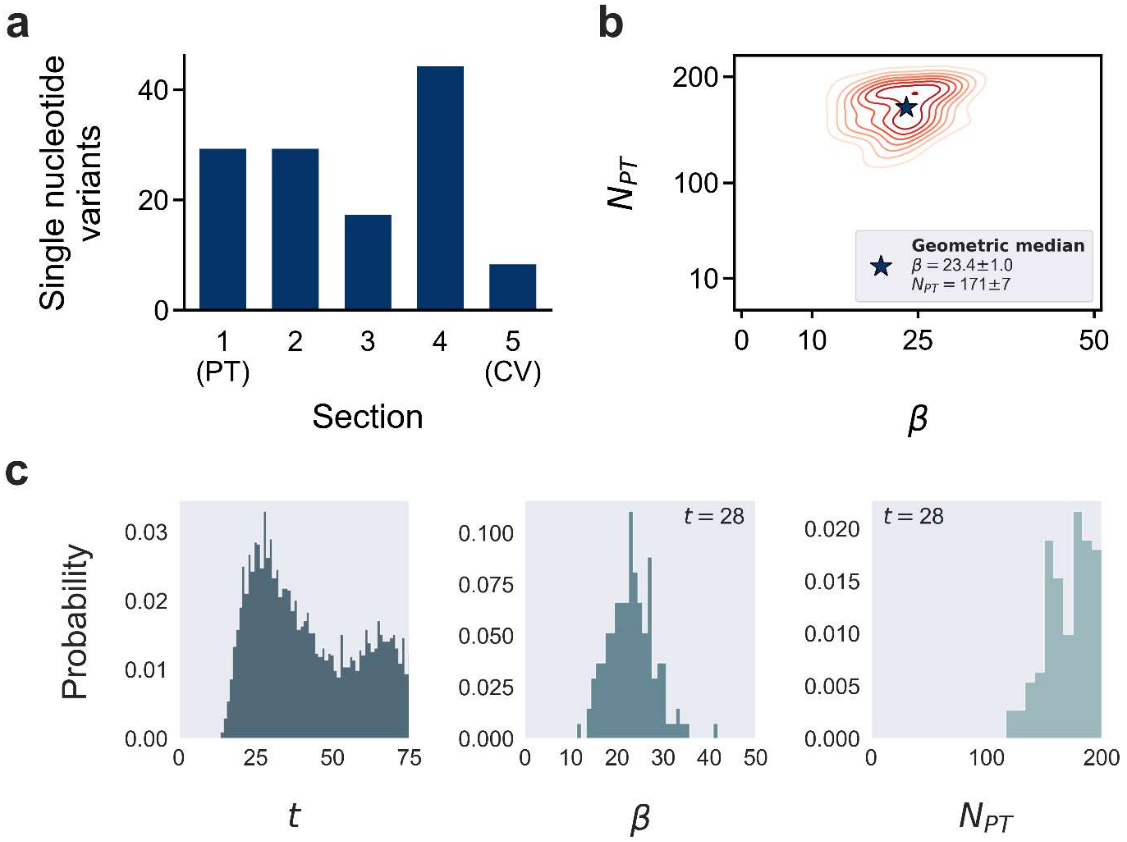
a
Single nucleotide variants
0
20
40
1
(PT)
2
3
4
5
(CV)
Section
b
$N_{PT}$
200
100
10
0
10
25
50
$\beta$
Geometric median
$\beta = 23.4 \pm 1.0$
$N_{PT} = 171 \pm 7$
c
Probability
0.03
0.02
0.01
0.00
0
25
50
75
$t$
$t = 28$
0.100
0.075
0.050
0.025
0.000
0
10
20
30
40
50
$\beta$
$t = 28$
0.020
0.015
0.010
0.005
0.000
0
100
200
$N_{PT}$

# 193B P11L2

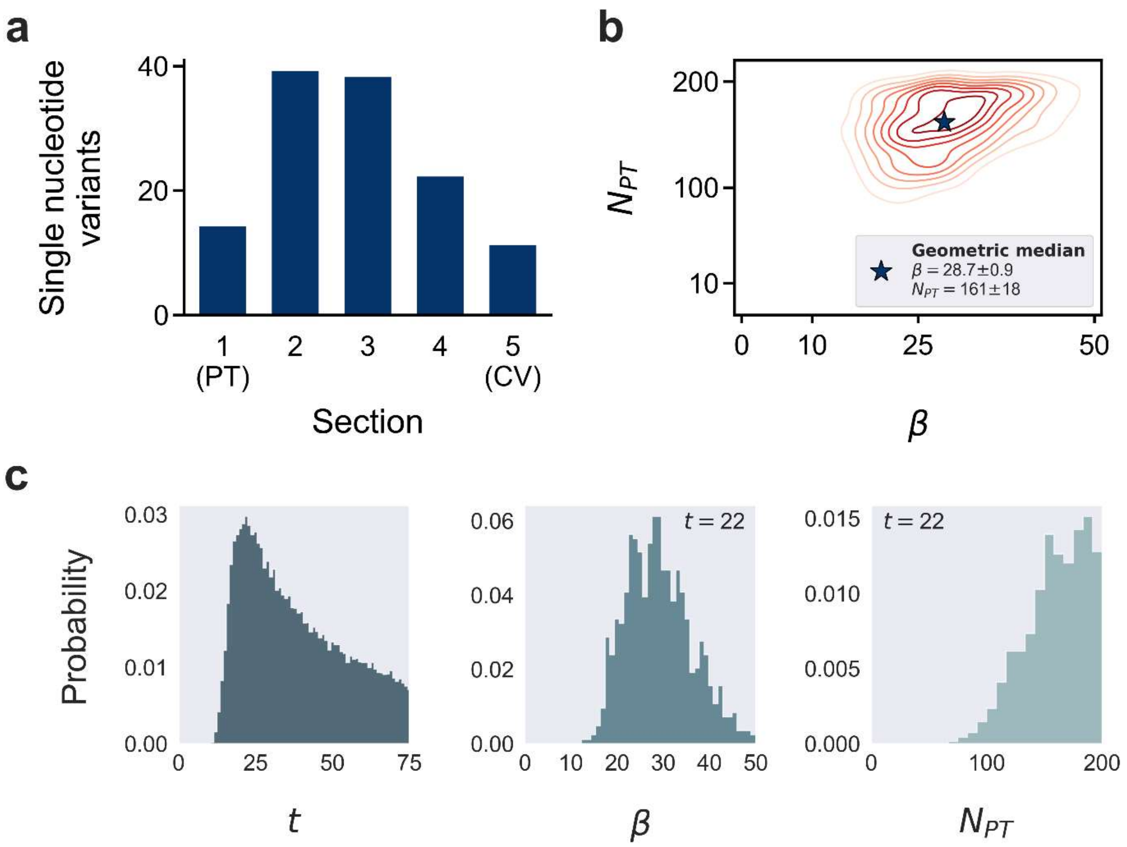
a
Single nucleotide variants
40
20
0
1 (PT)
2
3
4
5 (CV)
Section
b
$N_{PT}$
200
100
10
0
10
25
50
$\beta$
Geometric median
$\beta = 28.7 \pm 0.9$
$N_{PT} = 161 \pm 18$
c
Probability
0.03
0.02
0.01
0.00
0
25
50
75
$t$
$t = 22$
0.06
0.04
0.02
0.00
0
10
20
30
40
50
$\beta$
$t = 22$
0.015
0.010
0.005
0.000
0
100
200
$N_{PT}$

# 193B P11L1

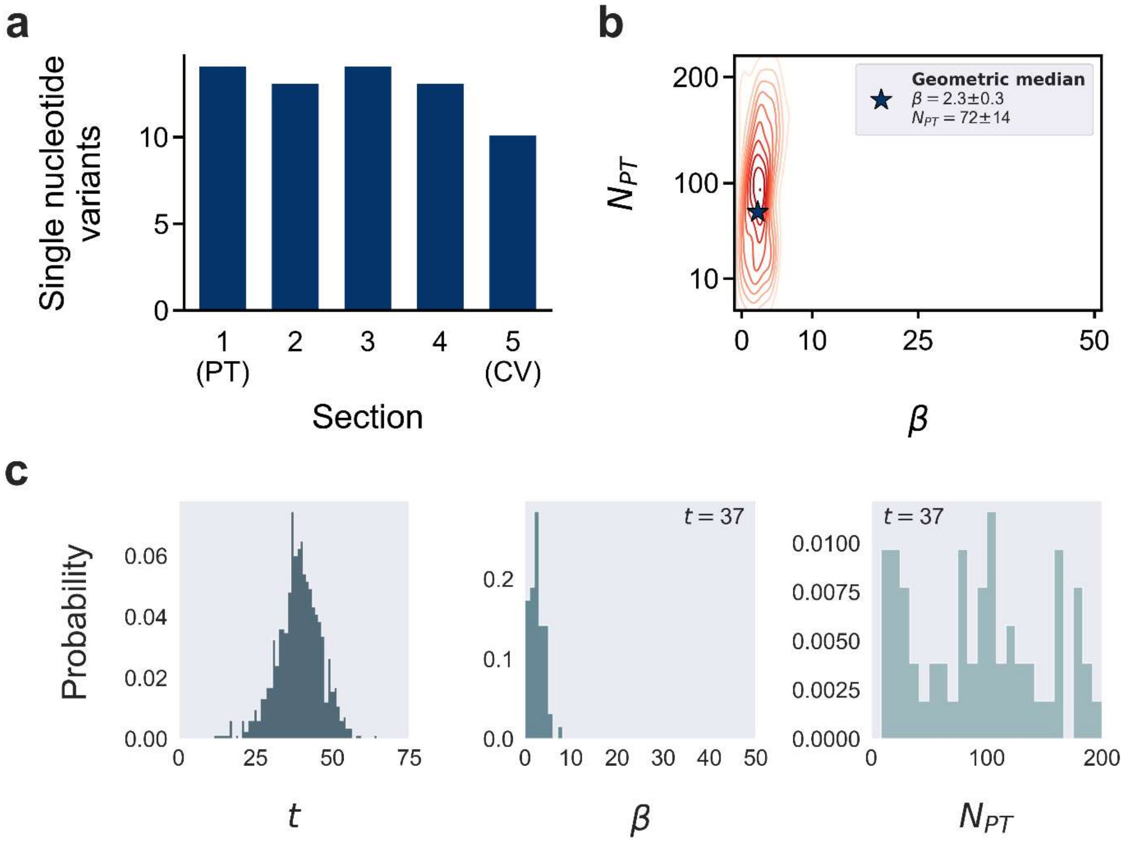
a
Single nucleotide variants
0
5
10
1 (PT)
2
3
4
5 (CV)
Section
b
Geometric median
β = 2.3±0.3
N_PT = 72±14
N_PT
200
100
10
0
10
25
50
β
c
Probability
0.00
0.02
0.04
0.06
0
25
50
75
t
t = 37
0.0
0.1
0.2
0
10
20
30
40
50
β
t = 37
0.0000
0.0025
0.0050
0.0075
0.0100
0
100
200
N_PT

# 193B P10L2

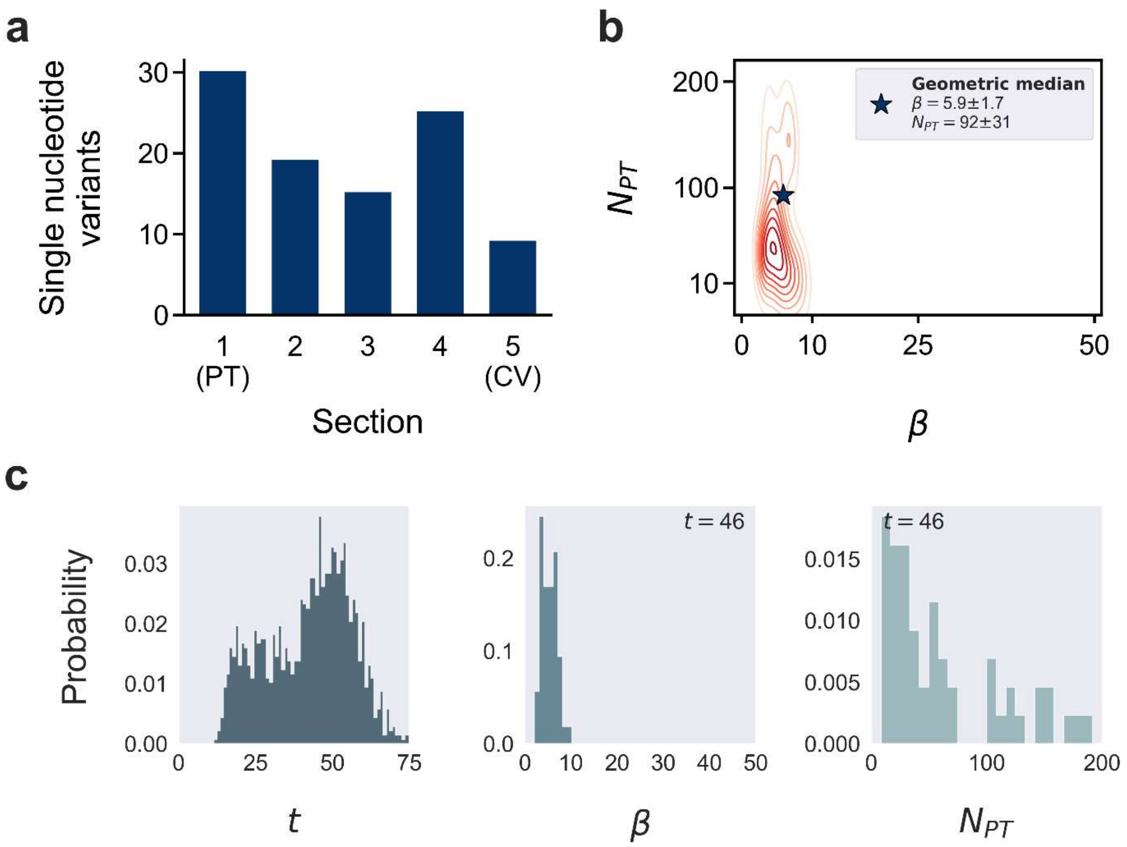
a
b
c
Single nucleotide variants
30
20
10
0
1
(PT)
2
3
4
5
(CV)
Section
Geometric median
β = 5.9±1.7
N_PT = 92±31
N_PT
200
100
10
0
10
25
50
β
Probability
0.03
0.02
0.01
0.00
0
25
50
75
t
t = 46
0.2
0.1
0.0
0
10
20
30
40
50
β
t = 46
0.015
0.010
0.005
0.000
0
100
200
N_PT

# 193B P10L1

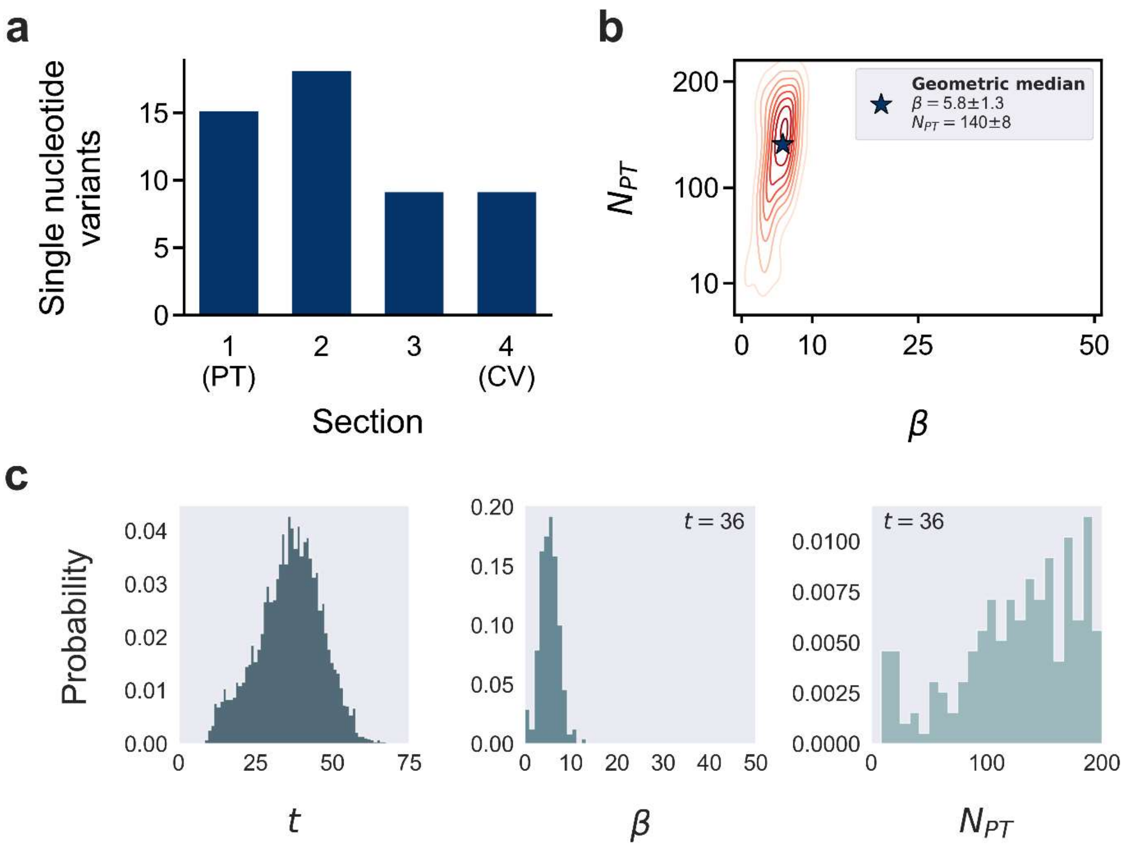
a
Single nucleotide variants
0
5
10
15
1
(PT)
2
3
4
(CV)
Section
b
$N_{PT}$
10
100
200
0
10
25
50
$\beta$
Geometric median
$\beta = 5.8 \pm 1.3$
$N_{PT} = 140 \pm 8$
c
Probability
0.00
0.01
0.02
0.03
0.04
0
25
50
75
$t$
0.00
0.05
0.10
0.15
0.20
$t = 36$
0
10
20
30
40
50
$\beta$
0.0000
0.0025
0.0050
0.0075
0.0100
$t = 36$
0
100
200
$N_{PT}$

# 193B P4L3

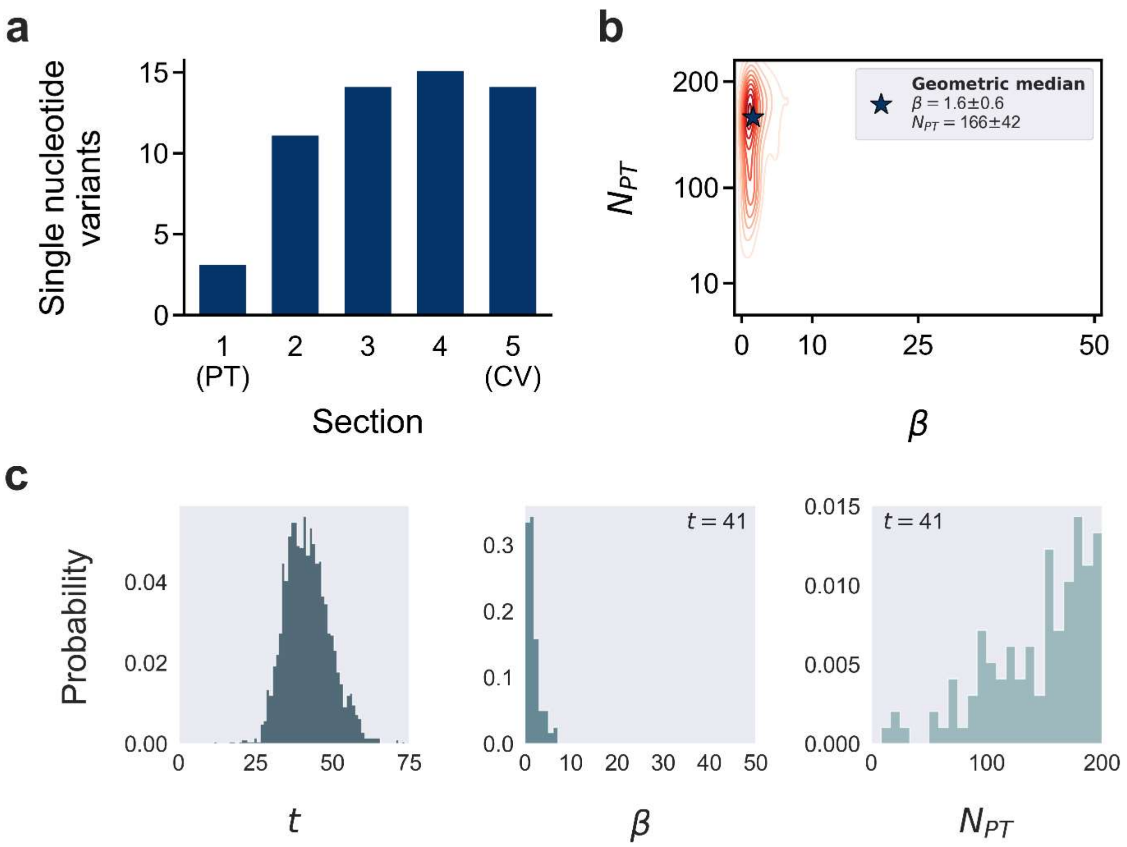
a
Single nucleotide variants
15
10
5
0
1 (PT)
2
3
4
5 (CV)
Section
b
Geometric median
β = 1.6±0.6
N_PT = 166±42
N_PT
200
100
10
0
10
25
50
β
c
Probability
0.04
0.02
0.00
0
25
50
75
t
t = 41
0.3
0.2
0.1
0.0
0
10
20
30
40
50
β
0.015
t = 41
0.010
0.005
0.000
0
100
200
N_PT

# 193B P4L1

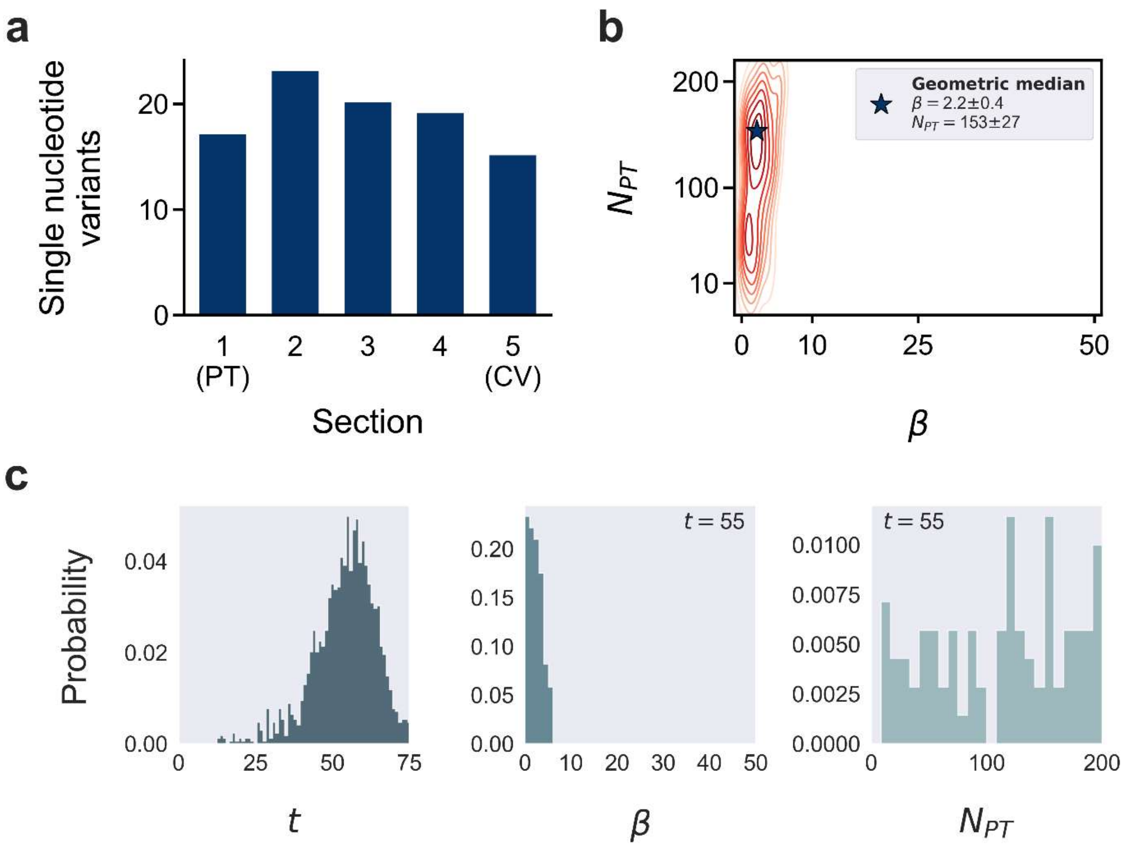
a
Single nucleotide variants
0
10
20
1 (PT)
2
3
4
5 (CV)
Section
b
Geometric median
$\beta = 2.2 \pm 0.4$
$N_{PT} = 153 \pm 27$
$N_{PT}$
10
100
200
0
10
25
50
$\beta$
c
Probability
0.00
0.02
0.04
0
25
50
75
$t$
$t = 55$
0.00
0.05
0.10
0.15
0.20
0
10
20
30
40
50
$\beta$
$t = 55$
0.0000
0.0025
0.0050
0.0075
0.0100
0
100
200
$N_{PT}$
